# Histone H3.3 ensures cell proliferation and genomic stability during myeloid cell development

**DOI:** 10.1101/2024.12.26.630087

**Authors:** Sakshi Chauhan, Anup Dey, Todd Macfarlan, Keiko Ozato

## Abstract

Variant histone H3.3 is thought to be critical for survival of many cells, since it is deposited in expressed genes, a feature different from core histones. For example, H3.3 deletion leads to embryonic lethality in mice. However, requirement of H3.3 in later stage of development has remained unclear. The aim of this work was to elucidate the role of H3.3 for development of myeloid lineage, important for innate immunity. We conditionally knocked out (cKO) the H3.3 genes in myeloid progenitor cells differentiating into bone marrow derived macrophages (BMDMs). Progenitor cells lacking H3.3 were defective in replication, suffered from extensive DNA damage, and underwent apoptosis. Surviving *H3.3*cKO cells expressed many interferon stimulated genes (ISGs) throughout differentiation. Further, *H3.3*cKO BMDMs possessed chromatin accessible sites, and histone posttranslational modifications consistent with the gene expression profiles, Accordingly, *H3.3*cKO BMDMs retained general nucleosomal structure genome wide. In summary, H3.3 is required for proliferation of myeloid progenitor cells, but is in large part dispensable for differentiation of BMDMs.

---

Histone H3.3, encoded by two genes, represents a variant deposited on nucleosomes of expressed genes throughout cell cycle. This feature is distinct from that of core histones, H3.1 and H3.2 that are deposited during replication. Because of transcription coupled deposition, H3.3 is implicated for epigenetic memory of expressed genes [1]. In addition to actively expressed genes, H3.3 is found in the heterochromatin and telomeres [2, 3].

H3.3 is essential for embryonic development, since deletion of H3.3 genes leads to early embryonic lethality, attributed to impaired proliferation[4].

Requirement of H3.3 for later stages of development, however, appears more complex. In some models, depletion of H3.3 genes does not totally abrogate developmental progression, while it alters direction of differentiation. For example, H3.3 deletion in hematopoietic stem cells is reported to result in myeloid lineage biased progenitor cells, displaying altered histone modification patterns [5]. Deletion of H3.3 in neuronal progenitor cells is shown to reduce the progenitor population but does not completely prevent generation of postmitotic neurons [6]. Recently it has been reported that H3.3 is downregulated during B lymphocytes development, and this process is necessary for plasma cells differentiation [7]. Limited requirement of H3.3 for differentiation in some systems is surprising, considering that H3.3 has a nonredundant role in being deposited in expressed genes.

In this study, we investigated the role of H3.3 in myeloid cell development. The myeloid lineage originates from HSCs, that progress to myeloid progenitor cells, which then differentiate into terminally differentiated monocytes/ macrophages. Myeloid lineage cells confer innate protection against pathogens and elicit inflammation, the role essential for life. We constructed mice in which both *H3f3a* and *H3f3b* genes were floxed, conditionally knocked out both genes in myeloid progenitors in bone marrow, using LysM-Cre. We investigated how H3.3 deletion affects development into bone marrow derived macrophages (BMDMs) in vitro. We chose the in vitro model, since it eliminates the potential effect of incoming and outgoing cell population, which is unavoidable with in vivo models.

We show that *H3.3*cKO progenitor cells succumb to DNA damage and undergo apoptosis. Accordingly, fewer BMDMs were generated from *H3.3*cKO progenitors. We found that *H3.3*cKO BMDMs expressed many interferon stimulated genes (ISGs), likely due to DNA damage. The ISG induction was mediated by canonical interferon pathway, but did not depend on STING, IRF7, or RIGI. Interestingly, *H3.3*cKO BMDMs possessed basic nucleosomal arrays, comparable to wild type BMDMs. *H3.3*cKO BMDMs also displayed chromatin accessible sites and posttranslational histone modifications concordant with the gene expression profiles. Together, H3.3 is obligatory for proliferation myeloid progenitor cells, but not completely required for differentiation into postmitotic BMDMs.

## Results

### Cell cycle delay and apoptosis in *H3.3*cKO cells

Histone H3.3 is encoded by two separate genes,- *H3f3a* on chromosome1 and *H3f3b* on chromosome 11. Each of these genes has 4 exons. To create *H3.3 ^fl/fl^* mouse, Loxp sites were inserted on flanking exons of *H3f3a* (exon 2) gene (Supplementary Fig 1a) and *H3f3b* (exon 1-4) gene (Supplementary Fig 1b) by homologous recombination. To create *LysM^cre/cre^* conditional KO, *H3f3a ^fl/fl^:H3f3b ^fl/fl^*mice were crossed with *LysM^cre/cre^* mice.

At genomic level, H3f3a and *H3f3b* deletion by *LysM^cre/cre^*was confirmed by PCR screening (Supplementary Fig1c). Depletion of H3.3 mRNA and protein in BMDMs was confirmed with the qRT-PCR (Supplementary Fig1d) and western blot analysis (Supplementary Fig1e) respectively.

We performed flow cytometry analysis of bone marrow derived macrophages (BMDMs) from WT and *H3.3*cKO mice using macrophage cell surface markers CD11b and F4/80. The number of *H3.3*cKO BMDMs were significantly lower, less than half of WT counterpart (Fig1a), which pointed to either proliferation defect and or increased apoptosis in *H3.3*cKO cells. Bone marrow cells, particularly progenitor cells, proliferate at early stages of culture and then differentiate into postmitotic BMDMs by day 7. CD11b and F4/80 expression profiles on day3, 5 and 7 cells are shown in Supplementary Fig2a. Both *H3.3* genes were deleted early, day3 and day5 (Supplementary Fig 2 b, c). As for peritoneal macrophages, we found no significant difference in their number between WT and H3.3cKO (Supplementary Fig1g), This was expected as macrophages are continuously repopulated in vivo.

**Fig 1.**
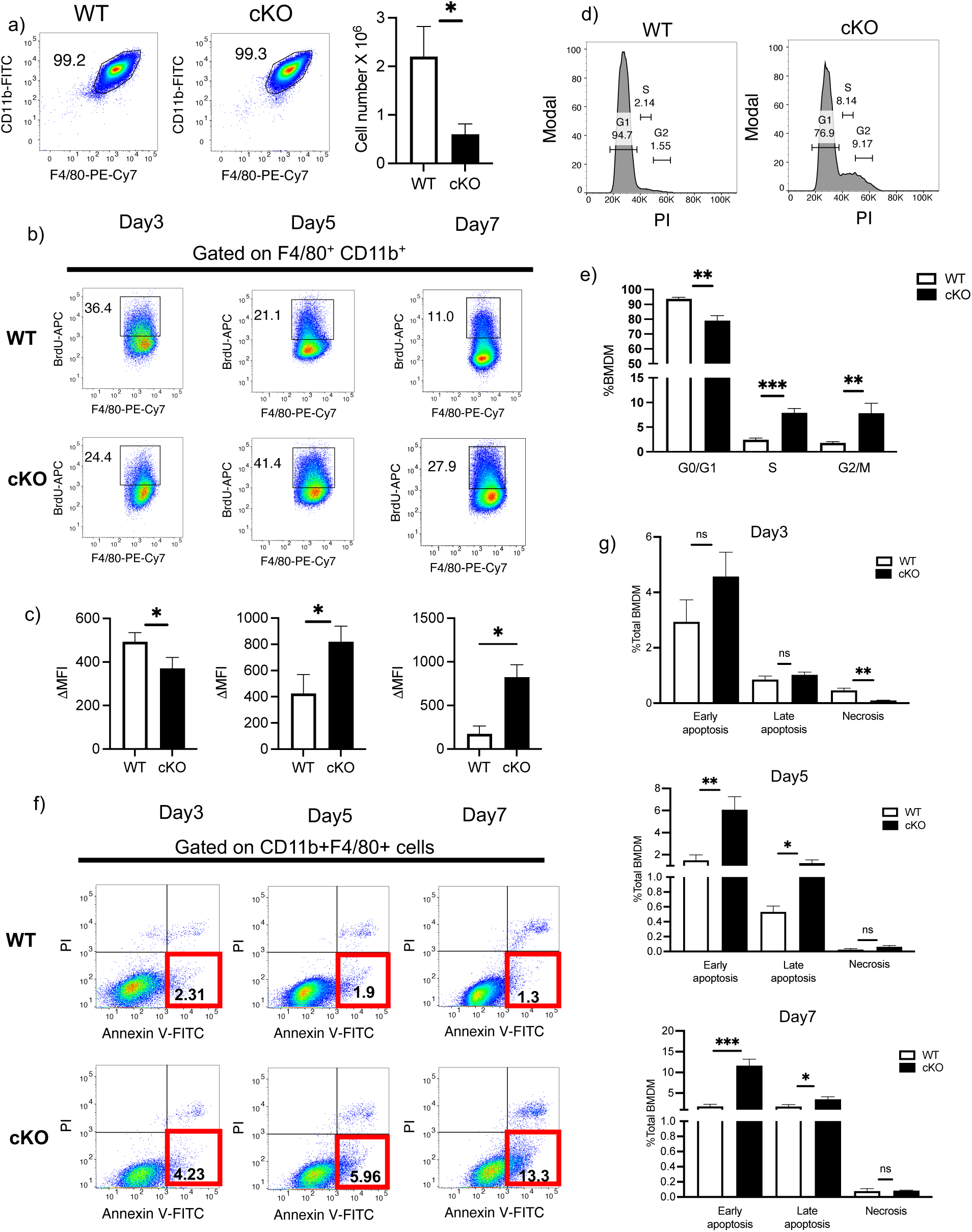
Proliferation defect and apoptosis in *H3.3*cKO BMDMs. a) Representative flow cytometry plots of bone marrow derived macrophages upon staining with CD11b and F4/80 markers. Total number of BMDMs (right) obtained from WT and cKO BM cells. b) Day3,5 and day7 cells were incubated with BrdU and subsequently analyzed by flow cytometry. c) Mean fluorescence intensities of BrdU were calculated for each day. d) Flow cytometric analysis of cell cycle analysis in PI stained BMDMs (day7). e) Percentage of cells were calculated for each phase of cell cycle. f) Annexin V-PI staining of WT and cKO cells was analyzed by flow cytometry to detect apoptosis. g) Cell percentage was calculated for early (Annexin V+, PI-), late (Annexin V+, PI+ and necrotic (Annexin V-, PI+) cells on day3, 5 and day7. a-g) n=3 mice. Unpaired t test was used to calculate P value. Bar graphs represent mean values +/- SD. NS - Not significant

To investigate cell cycle regulation in *H3.3*cKO cells, we performed BrdU incorporation assay for WT and *H3.3*cKO cells on day 3, day 5 and day 7. This assay identifies cells undergoing DNA replication. On day 3, BrdU uptake was significantly lower in *H3.3*cKO cells than WT cells, indicating that *H3.3*cKO cells were deficient in timely DNA synthesis (Fig 1 b,c). In line with reduced BrdU uptake, expression of Ki67, a proliferation marker was less in H3.3cKO cells than WT cells on day3 (Supplementary Fig3a-d). However, *H3.3*cKO cells had higher BrdU uptake on Day 5 and 7, indicating that untimely DNA synthesis continued in some of H3.3cKO cells (Supplementary Fig3a-d). In agreement with the result, propidium iodide staining revealed a small percentage of *H3.3*cKO cells were in S or G2/M phase, while most cells were in G1 (Fig 1d,e). These data show that in the absence of H3.3, bone marrow progenitor cells do not proliferate as well as WT cells, leading to the reduced BMDM yield.

It is possible that cells with defective proliferation may not survive and undergo apoptosis. To test this possibility, we performed Annexin V - PI staining. As shown in flow cytometry data in Fig1 (f, g), *H3.3*cKO cells exhibited higher Annexin V staining than WT cells. We also found that caspase 9, involved in apoptosis was cleaved in *H3.3*cKO cells (Extended Fig 3e) [8]. We also found that Poly (ADP-ribose) polymerase 1 (PARP-1) was cleaved in *H3.3*cKO cells, but not in WT cells (Supplementary Fig3e) [9]. These data led us to conclude that *H3.3*cKO cells are deficient in DNA replication and a certain fraction of cells die due to apoptosis.

### *H3.3*cKO cells succumb to DNA damage

We surmised that above observed apoptosis is a consequence of DNA damage. Indeed, we found that ATM and ATR were phosphorylated In *H3.3*cKO cells (Fig2a, b). ATM and ATR are kinases, activated upon dsDNA breaks, which then initiate DNA damage response pathway [10, 11], H2AX, a variant of histone H2A, is an immediate target of ATR, ATM. Phosphorylated H2AX, γH2AX is one of most widely used marker for DNA damage [12]. γH2AX is thought to bind to DNA damage sites, initiating DNA damage response (DDR) [13–15]. p53 is a transcription factor that activates many downstream genes to regulate apoptosis and cell cycle arrest such as p21 [16, 17].

**Fig 2.**
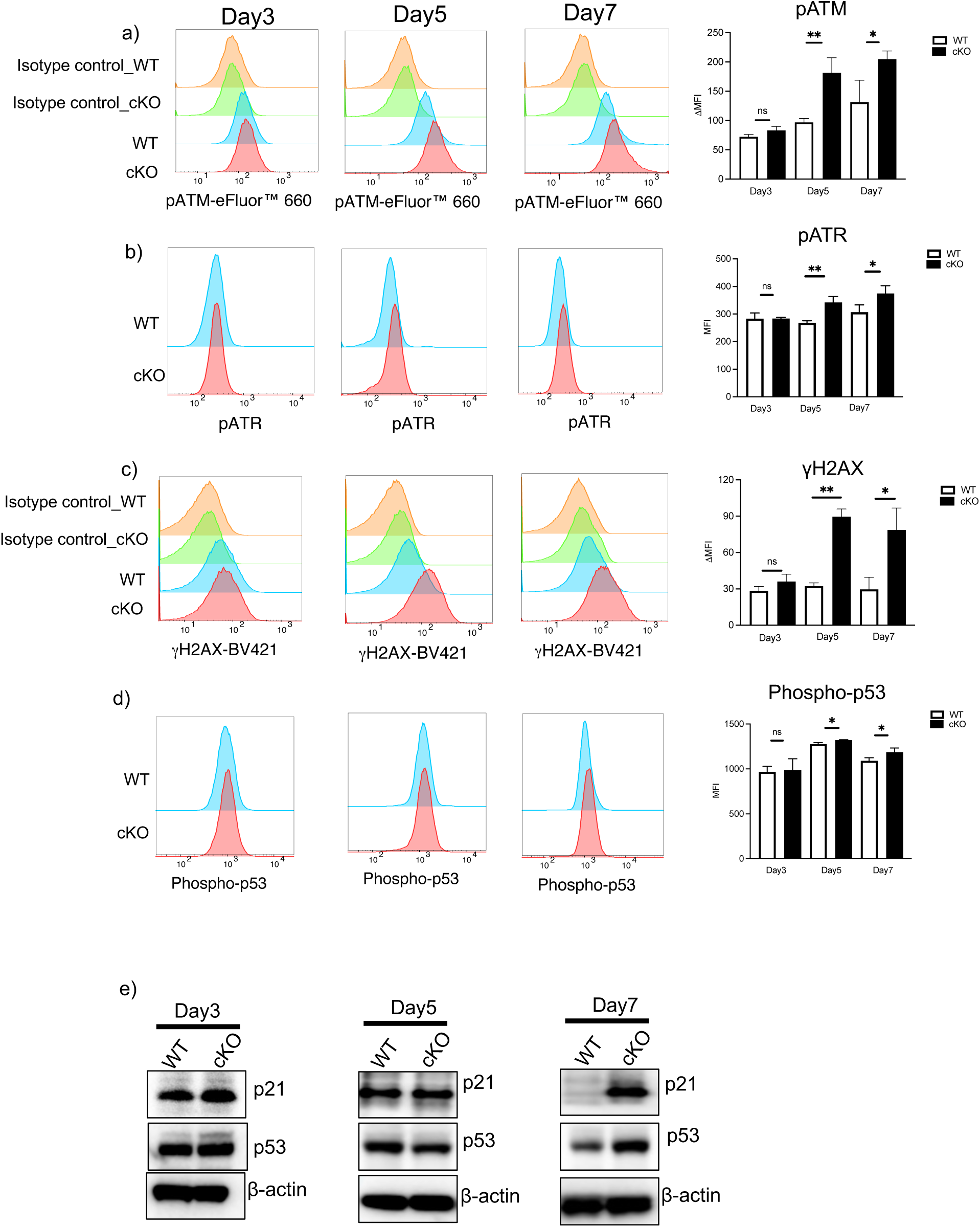
DNA damage in *H3.3*cKO BMDMs. a) Flow cytometric analysis and quantification of phosphorylated ATM and b) phosphorylated ATR c) phosphorylated H2AX d) phosphorylated p53 on day3, day5 and day7 BMDMs. MFI values represent the average of three WT and three H3.3 cKO mice ± SD. Unpaired t test was used to calculate P values. e) Immunoblot detection of p53 and p21 protein in WT and cKO cells on day 3,5 and 7.

Flow cytometry data in Fig 2a,b,c,d demonstrated phosphorylation of ATM, ATR, H2AX, and p53 in *H3.3*cKO cells on day 3, day 5 and day 7 (quantification on the right). Western blot data in Fig 2e confirmed overexpression of p21 in *H3.3*cKO cells on day 7, but not in WT cells. Moreover, we have previously shown that DNA damage occurs when cell cycle progression is inhibited [18]. In sum, *H3.3*cKO cells sustain DNA damage and harbor many factors involved in DDR.

### Nucleotide synthesis genes are downregulated in *H3.3*cKO cells

To delineate a mechanism of DNA damage in *H3.3*cKO cells, we undertook unbiased transcriptome profiling. RNA-seq was performed for *H3.3*cKO and WT cells on day3, day 5 and day 7 (later, on day 8) (Experimental Scheme in Fig 3a). On day 3, there were 1210 differentially expressed genes (upregulated in cKO - 781, downregulated in cKO - 429, FDR<0.05). GO analysis of the day3 DEGs revealed downregulation of genes important for nucleotide synthesis and ribonucleotide biosynthetic process (Fig3b) in cKO cells. Nucleotide metabolism is crucial for cell growth, as it supports DNA and RNA synthesis [19]. Examples of genes in these categories include *Tk1*, *Impdh1*, *Adssl1*, *Uprt*, *Gmpr* and *Paics* which code for the enzymes involved in the nucleotide synthesis (Fig 3c) [20]. Genes associated with carbohydrate biosynthesis, amino acid metabolism and cytoskeletal organization were also downregulated (Fig 3b). GO terms for upregulated genes included apoptosis regulated genes (Fig 3b). These data indicate that H3.3 is required for replication and cell cycle progression, and in its absence apoptotic genes are activated. These results are consistent with the phenotypic data described above.

**Fig 3.**
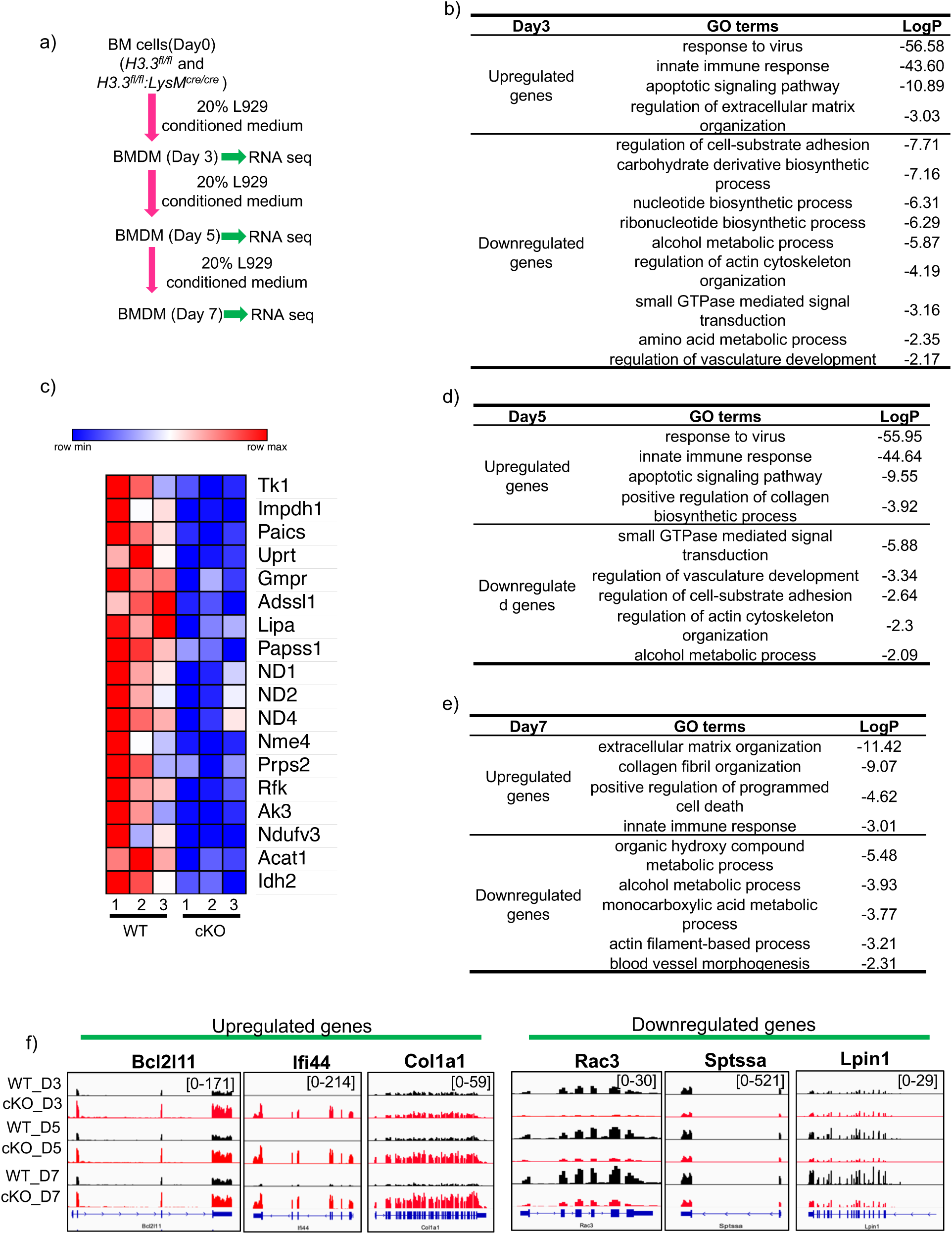
Transcriptome analysis of WT and cKO cells at different time points i.e. day3, 5 and 7. a) Schematic of RNA seq with day3, day5 and day7 BMDMs b) GO analysis of differentially expressed genes on day3. c) Heatmap shows downregulation of nucleotide/ribonucleotide biosynthesis genes like thymidine kinase 1(*Tk1*), adenylosuccinate synthase 1(*Adssl1*), inosine monophosphate dehydrogenase 1 (*Impdh1*), *Paics*, *Uprt* (uracil phosphoribosyl transferase) and *Gmpr* (guanosine monophosphate reductase) etc. on day3. d) GO analysis of cKO up and downregulated genes on day5 and e) on day7 f) IGV gene tracks of upregulated genes (apoptotic gene *Bcl2l11*, ISG *Ifi44* and collagen gene *Col1a1*) and downregulated genes (small GTPase *Rac3*, sphingolipid biosynthesis gene *Sptssa* and lipid synthesis gene *Lpin1*) show continuous up and downregulation respectively from day3 to day7.

RNA-seq on day 5 and 7 BMDMs revealed 638 (upregulated in cKO - 498, down regulated in cKO - 140, FDR<0.05) and 627 (upregulated in cKO - 372, downregulated in cKO - 255, FDR<0.05) DEGs, respectively.

GO analysis of day 5 and 7 did not reveal nucleotide biosynthetic process and ribonucleotide biosynthetic process as the most prominent GO categories, suggesting that genes in these categories may be more sensitive to H3.3 deletion in early stages of bone marrow cell development (Fig 3 d,e).

On the other hand, for upregulated genes, GO terms representing immune responses and response to viruses were found in *H3.3*cKO samples on day 3, 5 and day 7 (Fig 3 b,d,e). IGV examples of upregulated (*Bcl2l11, Ifi44, Col1*) and downregulated (*Rac3,Sptssa* and *Lpin1*) genes are presented in Fig 3f.

### Interferon stimulated genes (ISGs) are upregulated in *H3.3*cKO BMDMs

It was striking that genes representing innate immune responses and response to virus were upregulated in *H3.3*cKO cells at all stages of culture, from day 3 up to day 7. RNA-seq was also performed for BMDMs on day 8, which revealed 554 upregulated genes, while 309 gene were down regulated in *H3.3*cKO cells (FDR<0.05) (Fig4a). Out of 554 upregulated genes in cKO cells, many (311) were authentic ISGs, in that they were also induced after interferon-γ treatment. This is illustrated in Volcano plots of Fig 4a and Fig 4b. The former depicts all DEGs and the later ISGs only. Transcriptome analysis of *H3.3*cKO peritoneal macrophages also showed upregulation of innate immune response genes, mostly ISGs (Fig4 c,d). Fig 4 e,f,g shows enhanced mRNA and protein expression of *Stat1,* an ISG in *H3.3*cKO BMDMs and peritoneal macrophages respectively. Extensive ISG expression would indicate that *H3.3*cKO BMDMs might be under inflammatory state, given many ISGs encode inflammation inducing cytokines and chemokines.

**Fig 4.**
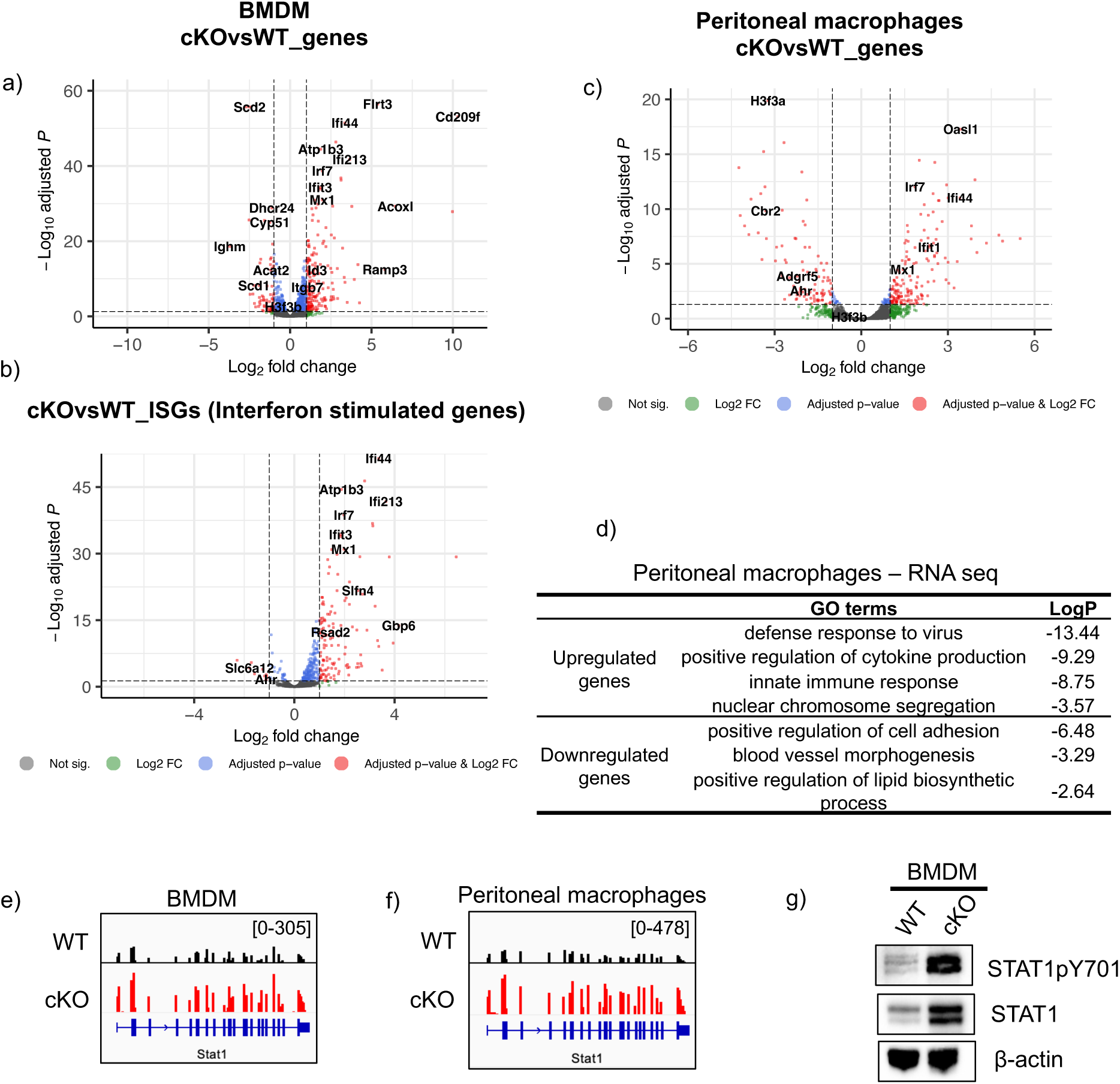
ISG expression in *H3.3*cKO BMDM and peritoneal macrophages. a) Volcano plots for differentially expressed genes and b) ISGs in bone marrow derived macrophages c) GO analysis of differentially expressed genes in bone marrow derived macrophages upon *H3.3* deletion (on day8 of differentiation). d) Volcano plot for differentially expressed genes in peritoneal macrophages e) GO analysis of differentially expressed genes in peritoneal macrophages upon *H3.3* deletion f) IGV browser screenshots showing *Stat1* gene upregulation in BMDM and g) peritoneal macrophages h) Western blot showing enhanced expression of Stat1 protein and its phosphorylated form in cKO BMDMs.

### STING, MAVS and IRF7 are dispensable for ISG induction in *H3.3*cKO BMDMs

We found ISG induction in *H3.3*cKO cells intriguing, since DNA damage has been reported to cause ISG expression in various cells [21–24].

In this scenario, small nucleotides produced by DNA damage would be sensed by pattern recognition receptors (PRRs), which initiates interferon expression. This would lead to full ISG expression by a second, feedback step through the engagement of type I interferon receptors (IFNAR) [25, 26].

To verify that ISG expression in *H3.3*cKO cells relies on canonical interferon induction pathways, we examined whether neutralization of IFNAR1 could inhibits ISG expression in *H3.3*cKO cells. Results in Supplementary Fig 4a-d confirmed that blocking of IFNAR receptor resulted in marked reduction of ISG expression. Thus, ISG expression in *H3.3*cKO cells occurs through a canonical interferon feedback pathway.

We then sought to find the mechanism that initiates the first interferon expression in *H3.3*cKO cells. Small nucleotides, endogenous or exogenous are sensed by factors in the STING and RIGI-MAVS pathways [27–29]. Both *Sting* and *RigI*, themselves ISGs, were found upregulated in *H3.3*cKO cells (Fig 5a) To determine if one of these pathways plays a role, we constructed double knockout mice lacking STING and H3.3 (*H3.3^fl/^*^fl^:*LysM*^cre/cre^:*StingKO)* as well as those lacking MAVS and H3.3 (H3*.3^fl/^*^fl^:*LysM*^cre/cre^:*MavsKO*). RNA-seq was carried out for BMDMs generated from the double KO mice. Heat maps in Fig 5b showed that that deletion of *Sting* or *Mavs* did not abolish ISG expression in *H3.3*cKO cells. We thus concluded that STING, RIGI-MAVS pathways are not responsible for ISG induction in *H3.3*cKO cells. Finally, we tested if IRF7 plays a role in ISG expression in H3.3cKO cells. IRF7 is a transcription factor critical for host resistance as it activates many downstream ISGs [39]. IRF7 may also be activated by DNA damage [30, 31]. Double knockout mice lacking IRF7 and H3.3 (H3*.3^fl^* ^/fl^:*LysM*^cre/cre^:*Irf7KO*) were constructed, and RNA-seq was performed for BMDMs as above. Heat maps in Fig 5b again showed overall good ISG expression in double KO BMDMs, although levels of some ISGs were somewhat lower. Together, factor(s) that initiates ISG induction is in *H3.3*cKO cells remains to be identified.

**Fig 5.**
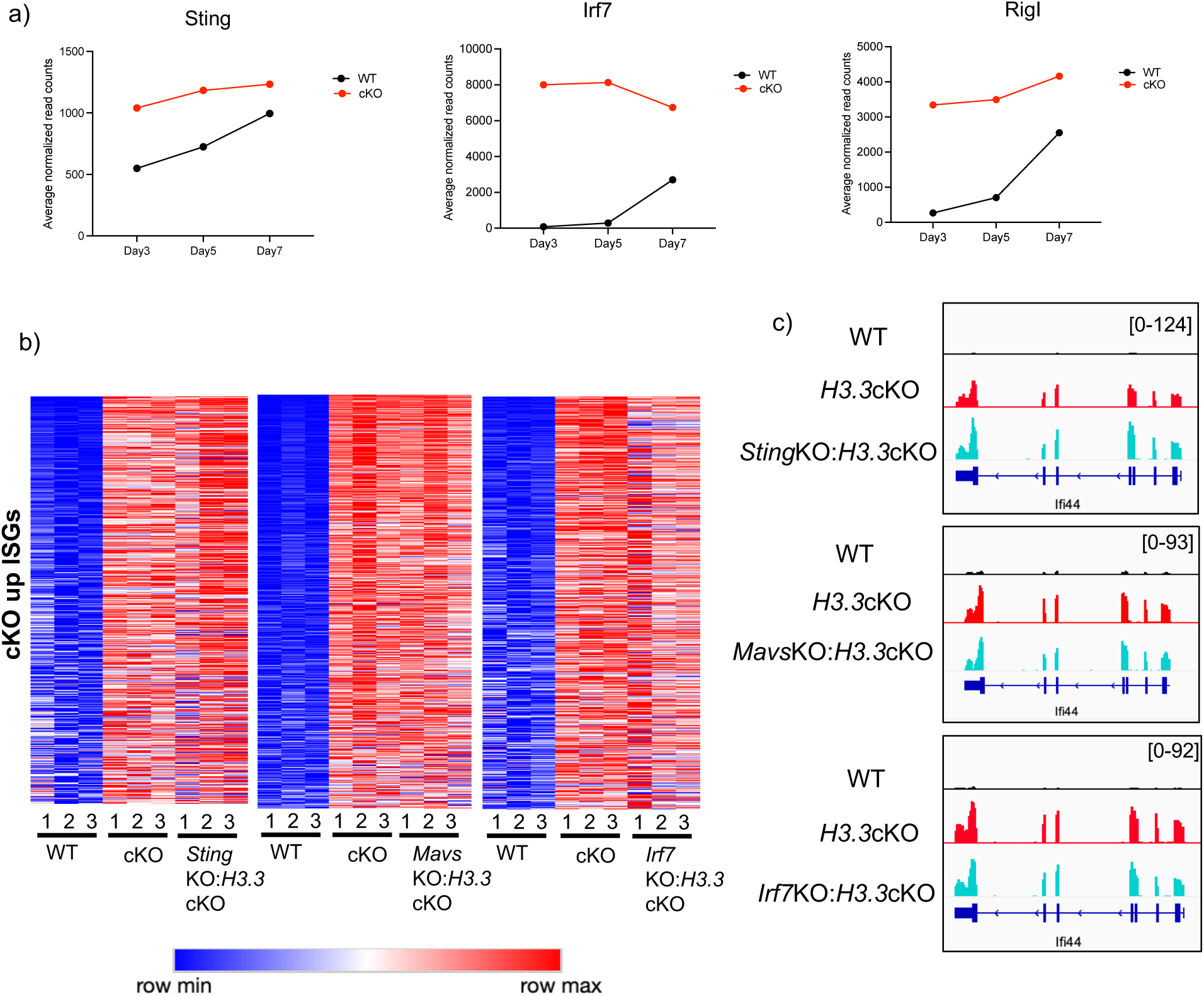
*Sting, Irf7* and *Mavs* are dispensable for antiviral response in cKO cells. a) Normalized read counts of *Sting*, *Irf7* and *RigI* in WT and cKO cells on day 3, 5 and 7. b) Heatmap shows expression of cKO upregulated ISGs is unperturbed in *H3.3*cKO: *Sting* KO, *H3.3*cKO: *Irf7* KO and *H3.3*cKO: *Mavs* KO BMDMs. c) IGV browser screenshots shows expression of *Ifi44* gene across WT, cKO, *H3.3*cKO: Sting KO, *H3.3*cKO: *Mavs* KO and *H3.3*cKO: *Irf7* KO BMDMs.

### H3.3 deletion does not completely prevent terminal differentiation of BMDMs

Bone marrow progenitor cells, upon several cycles of proliferation differentiate into postmitotic, terminally differentiated BMDMs. It was somewhat surprising that *H3.3*cKO gave rise to terminally differentiated BMDMs, although reduced in number. *H3.3*cKO BMDMs exhibited typical macrophage morphology, adhered to the plate like WT BMDMs (Supplementary Fig5a). Moreover, they expressed many genes specifying macrophages, although they expressed ISGs, unlike WT counterparts (Supplementary Fig 5b-e). In addition, *H3.3*cKO BMDMs retained the ability to respond interferonγ and increased ISG expression (Supplementary Fig 5f, g). Thus, the absence of H3.3 did not prevent bone marrow progenitor cells to develop into BMDMs, although it altered the transcriptome profiles to some degree. These results indicate that H3.3 is dispensable for BMDM terminal differentiation. Similar observation was reported with respect to postmitotic neurons [6], however, other cell types like ESCs require H3.3 for differentiation [32–35].

### H3.3 localizes to transcriptionally active chromatin

To identify genome-wide H3.3 deposition sites, CUT & RUN seq was carried out using BMDMs generated from knock-in mice in which H3.3 genes were replaced by HA-tagged H3.3 [36]. We found a total of 27645 H3.3-HA peaks and genomic distribution puts majority of the peaks on the gene bodies (57.09%) (Fig6a). H3.3 deposition was higher on expressed genes than silent genes (Fig6b, c, Supplementary fig 6a), consistent with transcription coupled incorporation. GO analysis revealed that H3.3 peaks are linked to immune regulation, representing biological function of BMDMs (Fig6d). Additionally, 30.46% of total H3.3 peaks were found in the intergenic regions, where H3.3 binding was enriched at presumed enhancer regions (Fig6e, see below). We also found some cell cycle genes occupied by H3.3, which may be attributed to active transcription of cell cycle genes in the progenitor stage (Fig6f, g, Supplementary Fig 6b).

**Fig 6.**
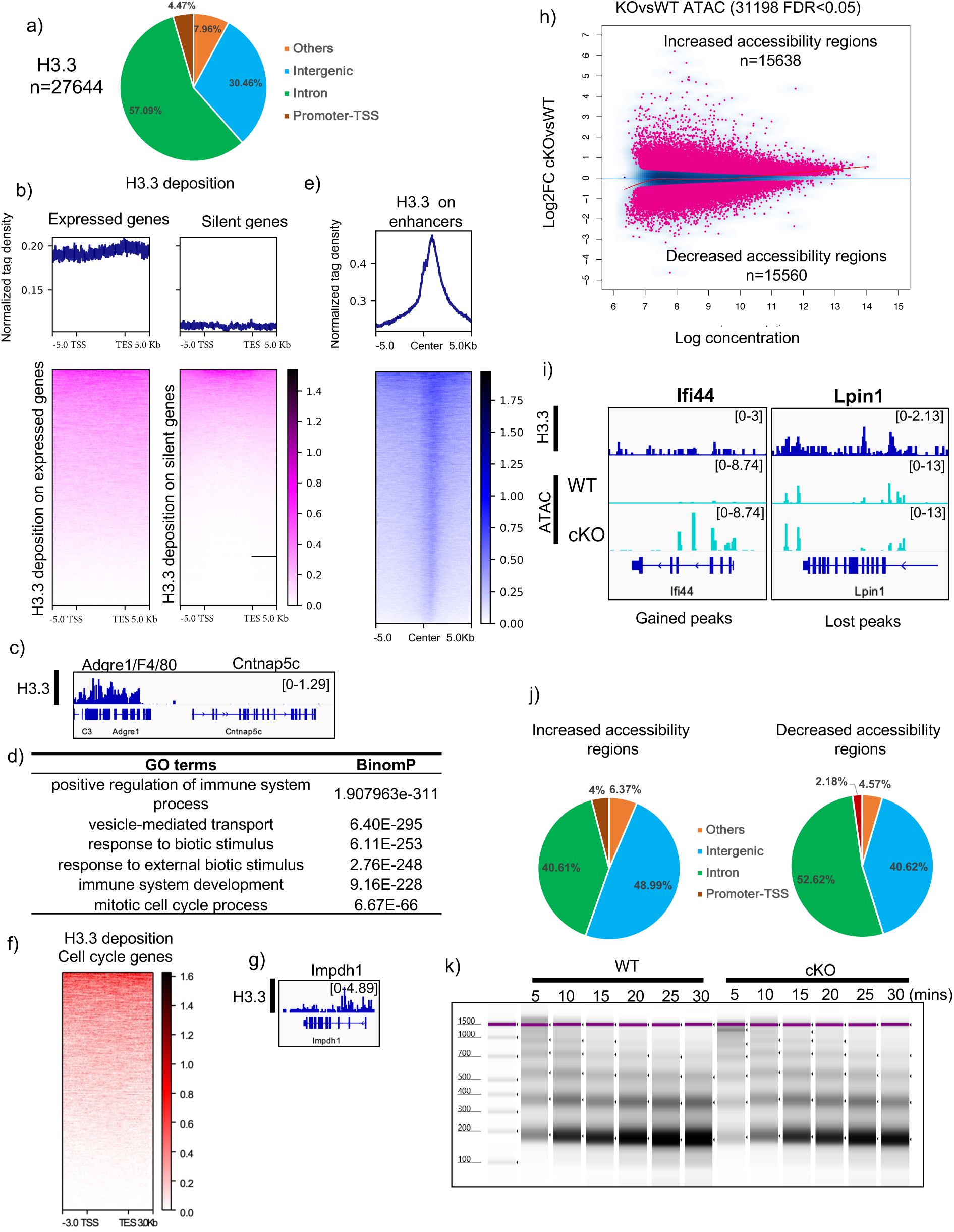
H3.3 distribution and chromatin accessibility analysis in cKO BMDMs. a) Pie chart shows H3.3 binding pattern in WT cells. b) Histogram and heatmap shows binding of H3.3 on expressed genes and on silent genes/isoforms. c) IGV tracks show deposition of H3.3 on expressed gene *Adgre1* and absence of H3.3 on silent gene *Cntnap5c*. d) GO analysis of H3.3 binding peaks. e) Profile plot and heatmap display deposition of H3.3 on enhancer regions. f) Heatmap shows H3.3 binding on cell cycle genes. g) IGV gene tracks depict H3.3 binding on a cell cycle gene (*Impdh1*). h) MA plot depicts increased and decreased accessible regions upon H3.3 deletion. i) Gene tracks show increased accessible peaks at *Ifi44* and decreased accessible peaks at *Lpin1* gene in cKO cells. H3.3 binding at these genes is shown in the top panel. j) Pie charts reveal genome wide distribution of cKO gained and lost peaks upon chromatin accessibility analysis. k) MNase mediated digestion of chromatin from WT and cKO day7 BMDMs.

### Chromatin accessibility patterns match transcriptome profiles of *H3.3*cKO BMDMs

We performed ATAC-seq, which reveal genomic sites open to transcription factors. We found 31198 differential sites (FDR <0.05) between WT and *H3.3*cKO BMDMs where *H3.3*cKO cells gained 15638 sites, while losing 15560 sites (Fig 6h). GO analysis revealed that the gained peaks were associated with innate immune response, cellular response to DNA damage stimulus and apoptosis. Lost peaks were linked to regulation of cell adhesion, regulation of leucocyte activation (Supplementary Fig 6c, d). Many of gained peaks were of the genes whose expression was gained in *H3.3*cKO, including ISGs. Examples of genes with gained or lost ATAC-seq peaks are presented in Fig 6i, namely *Ifi44* and *Lpin1*. As shown in Fig 6j, majority of gained peaks were deposited in the intergenic regions, while lost peaks were mostly in intronic region of H3.3cKO cells. De novo motif analysis revealed enrichment of the Atf3(bZIP), Sfpi motifs in the gained peaks, while Sfpi1 motifs were also present in the lost peaks (Supplementary Fig6e). These results indicate that *H3.3*cKO cells were able to create new accessible sites, while eliminating others. These changes likely provided a basis for transcriptome programs characteristic of *H3.3*cKO cells.

This observation prompted us to examine whether *H3.3*cKO cells maintain nucleosomal organizations without H3.3. We thus performed micrococcal nuclease (Mnase) digestion assay. As shown in Fig 6k, the digestion patterns were very similar between WT and *H3.3*cKO samples at all time points tested. These data indicate that *H3.3*cKO cells were able to form nucleosomal organizations without H3.3.

### Genes upregulated in *H3.3*cKO cells gain H3K27ac and H3K36me3 marks

Posttranslational histone modification marks (PTMs) that denote gene expression states are reported to be enriched on H3.3 [5, 37, 38]. We examined whether *H3.3* deletion results in changes in histone H3 modification marks, focusing on H3K27ac and H3K36me3. H3K27ac is a mark for active gene expression, enriched in enhancers of various sizes [39]. CUT & RUN experiments were carried out to determine genome-wide H3K27ac sites in WT and *H3.3*cKO BMDMs. In WT cells, majority of the H3K27ac peaks were found on gene bodies (Supplementary Fig7a, b). Differential binding analysis revealed that there were 4370 peaks with increased H3K27ac and 3022 peaks with reduced H3K27ac (FDR<0.05) in *H3.3*cKO cells (Fig7a, Extended Dat Fig7c). A significant number of gained and lost peaks were found in the intron. Whereas 16.35% of the gained peaks were in the promoter-TSS region (Fig7b). Volcano plot analysis revealed gained H3K27ac peaks were associated with genes involved in defense response to virus, programmed cell death and DNA damage regulation. On the other hand, response to sterol and blood vessel morphogenesis were associated with lost peaks (Supplementary Fig7d). For instance, *Ifi44,* an ISG expressed in *H3.3*cKO cells gained H3K27ac upon H3.3 deletion. Whereas *Lpin1* downregulated in *H3.3*cKO cells exhibited reduced H3K27ac peaks (Fig7d).

**Fig 7.**
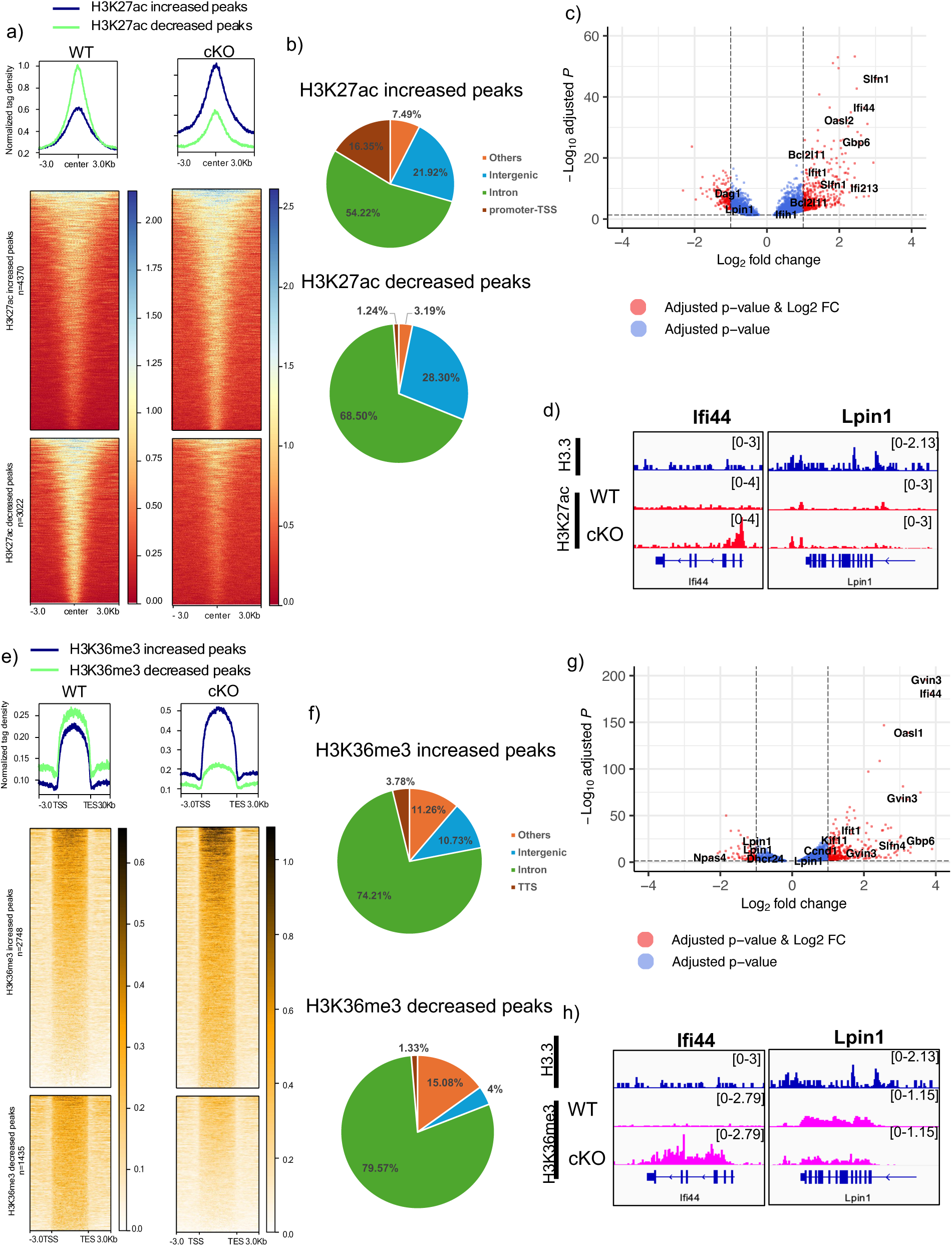
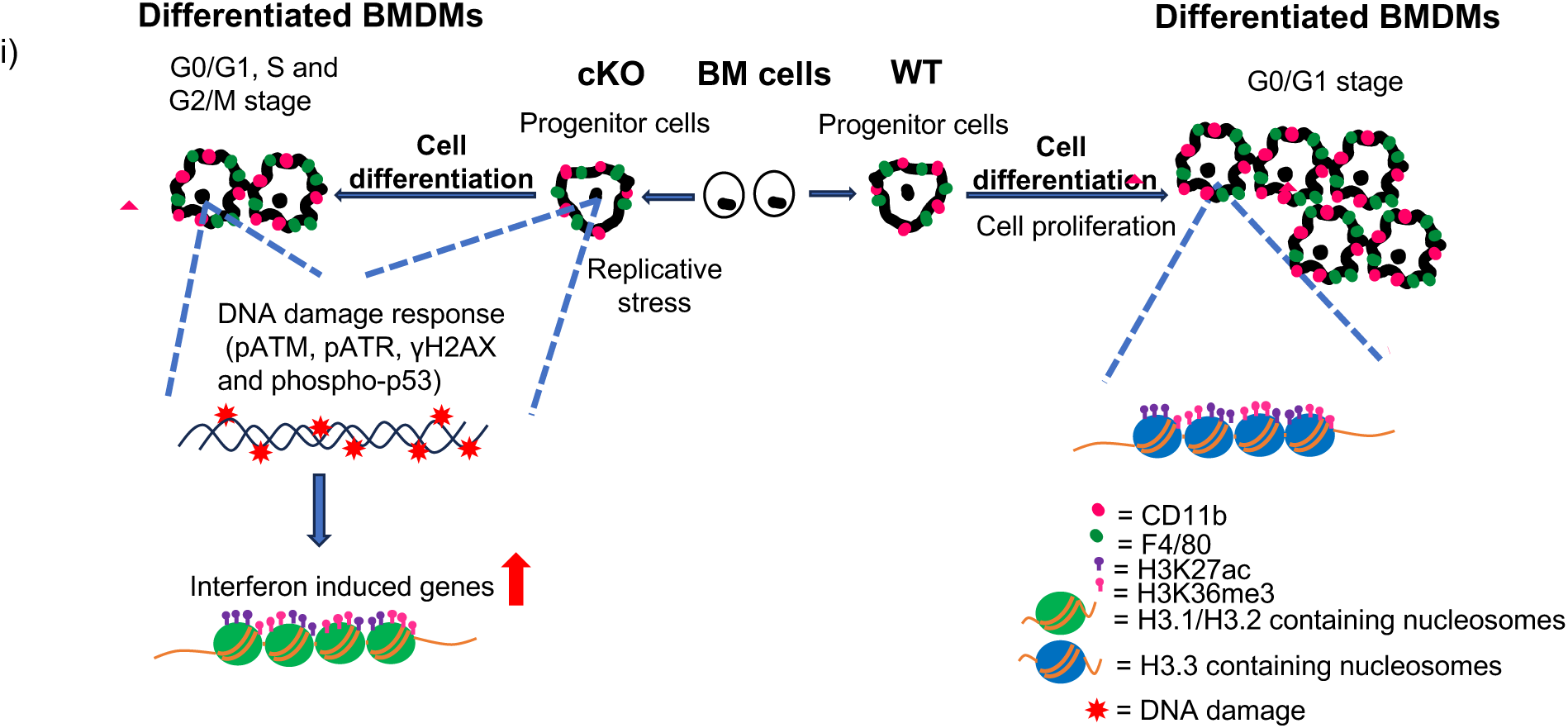
Differential distribution of H3K27c and H3K36me3 in cKO cells. a) Histograms and heatmaps of the H3K27ac densities for H3K27ac increased and decreased regions in WT and cKO cells. b) Pie charts represent genomic distribution of H3K27ac increased and decreased peaks. c) Volcano plot showing a correlation between H3K27ac CUT & RUN seq peaks and RNA seq data. The nearest genes for increased or decreased H3K27ac peaks correlate with genes up or downregulated in *H3.3*cKO BMDM. d) IGV tracks of genes associated with H3K27ac increased (*Ifi44* gene) or H3K27ac decreased peaks (*Lpin1*) show H3.3 and H3K27ac distribution in WT and cKO cells. e) Histograms and heatmaps of the H3K36me3 densities for H3K36me3 increased and decreased regions in WT and cKO cells. f) Genomic distribution of H3K36me3 increased and decreased peaks is shown in pie charts. g) Gained and lost peaks of H3K36me3 are shown on *Ifi44* and *Lpin1* genes respectively. Top panel shows H3.3 binding on these genes. h) IGV tracks for H3.3-HA (top) and H3k36me3 (middle two WT vs H3.3cKO) for *Ifi44 and Lpin1*. i) Model - BMDM development in *H3.3*cKO and WT mice <u>Right</u> (WT)): myeloid progenitors proliferate early on (day3) and produce many terminally differentiate into BMDMs by day7. <u>Left</u> (*H3.3*cKO): myeloid progenitors encounter replication stress and DNA damage and apoptosis, leading to fewer BMDMs. Surviving *H3.3*cKO cells express ISGs. *H3.3*cKO BMDMs form nucleosomal structure without H3.3 support.

The H3K36me3 mark has been linked with active transcription, DNA damage repair, and contribute to heterochromatin formation [40][41]. CUT & RUN seq for H3K36me3 revealed majority of the peaks on gene bodies in WT cells (Supplementary Fig7e, f). H3.3cKO cells had 2748 gained peaks and 1435 peaks H3K36me3 peaks for H3K26me3 (FDR<0.05) (Fig7e, Supplementary Fig 7g). Majority of gained and lost peaks were in the gene bodies (Fig7f). As expected, gained regions were found to be associated with response to interferon beta, chromosome organization and mitotic cell cycle GO terms, while lost peaks were lipid biosynthetic process and cholesterol metabolic process genes (Fig7g, Supplementary Fig7h). An example in Fig7h shows that *Ifi44* gained H3K36me3 marks in H3.3 cKO cells, while *Lpin1* lost the mark, similar to H3K27ac above. Also, genes having H3.3 deposition possessed H3K27ac and H3K36me3 marks (Supplementary Fig7i,). Moreover, genes upregulated in *H3.3*cKO cells gained H3K27ac as well as H3K36me3 marks, while those downregulated in *H3.3*cKO cells lost these modification marks. These results indicate that core histones H3.1 or H3.2, presumably substituting for H3.3,can gain PTMs associated with active gene expression.

## Discussion

The most salient findings we made in this study were (i) deletion of H3.3 leads to DNA damage and apoptosis in bone marrow progenitors, yielding fewer terminally differentiated BMDMs. (ii) *H3.3*cKO cells, both progenitor and BMDMs aberrantly expressed many ISGs, likely setting a prolonged inflammatory state. (iii) Deletion of H3.3, however, did not prevent BMDMs from retaining genome-wide chromatin structure and post-translational histone modifications.

DNA damage in *H3.3*cKO progenitor cells was mediated by the canonical DDR response, starting with the activation of ATM and ATR, followed by phosphorylation of H2AX (γH2AX) initiating assembly of DDR factors. In accordance, activated p53 was present in *H3.3*cKO cells, which led to expression of many downstream DDR genes, including p21 [42, 43]. In line with our findings, Jang et. al., reported that deletion of H3.3 from early embryos causes DNA damage, preventing further development [4]. It is possible that DNA damage ensues upon deletion of H3.3 in many cell types. However, DNA damage has not been documented in other H3.3 knockout models, including HSCs [5, 6, 44].

What is the mechanistic basis of DNA damage? Transcriptome data on day 3 showing that genes involved in nucleotide synthesis and ribonucleotide biosynthetic were down regulated in *H3.3*cKO cells strongly indicate that deletion of H3.3 interferes with DNA replication. *H3.3*cKO cells show reduced BrdU uptake on day 3 and abnormal uptake on day 5 and day 7 indicate that many of *H3.3*cKO cells are under replication stress. Given that replication stress is known to cause dsDNA breaks [45, 46], it seems safe to assume that defective replication accounts for DNA damage in *H3.3*cKO cells.

We observed that many ISGs were upregulated in *H3.3*cKO cells throughout, from day 3 to day7/8, progenitor to differentiated BMDMs. This ISG induction was attributed to the authentic JAK/STAT pathway. Observed ISG induction may be a consequence of DNA damage, since ISG expression has been documented upon DNA damage caused by drugs and pathogen infection [21–24]. To further understand ISG expression, we examined double KO mice lacking STING, MAVS, IRF7 which recognize endogenous nucleotides and amplify ISG induction, respectively [27, 28, 47]. We found none of these to be responsible for ISG induction in *H3.3*cKO cells. Thus, a sensing mechanism(s) other than those tested here seems to control ISG expression in *H3.3*cKO cells.

It is interesting to note that a considerable number of *H3.3*cKO cells survived DNA damage and apoptosis and differentiated into BMDMs. *H3.3*cKO BMDMs displayed morphological traits similar to WT BMDMs, and expressed many genes specific for macrophages, such as F4/80, CD11b. These results indicate that H3.3 is not totally required for generating terminally differentiated BMDMs, although the deletion reduced the number of BMDMs, and changed gene expression profiles, inducing ISGs. Supporting the view that H3.3 is not completely required for terminal differentiation, Funk et. al reported that deletion of H3.3 from neuronal progenitor cells did not prevent generation of postmitotic neurons, although inhibited proliferation [42]. In accordance with these findings, *H3.3*cKO BMDMs possessed nucleosomal arrays that were overall similar to those of WT cells (Fig 6). Consistent with chromatinized genome, *H3.3*cKO BMDM had DNA accessible sites whose numbers were comparable to WT cells. Furthermore, nucleosomes of H3.3cKO BMDMs were decorated with posttranslational histone modifications reflective of gene expression profiles (Model in Fig 7). It is possible that in the absence of H3.3, core histones H3.1 and/or H3.2 compensate, and incorporated into expressed genes allowing the maintenance of chromatinized genome. It is also possible that another variant histone H3, yet to be identified may be able to substitute for the lack of H3.3 [48]. In summary, this study highlights the critical requirement of H3.3 for DNA replication in the progenitor and development of the myeloid lineage.

## Materials and methods

### Mice

All mice used in this study were bred and maintained in the NICHD animal facility. All animal protocols utilized in this study, were approved by the Animal Care and Use Committee at National Institute of Child Health and Human Development (Animal Study Program #17-044, #20-044 and #23-044). Mice used in this study were derived from a C57BL/6 genetic background. H3.3^fl/fl^ mice were crossed with LysMcre mice (strain # 004781, The Jackson Laboratory). *Sting* KO [49] and *Irf7* KO [47] mice were crossed with *H3.3^fl/fl^;LysM^cre/cre^* mice to generate *StingKO:H3.3cKO* and *Irf7KO:H3.3cKO* mice respectively. *Mavs* KO mice (Strain # 008634, The Jackson Laboratory) were backcrossed to *H3.3^fl/fl^;LysM^cre/cre^*(C57BL/6 background) strain over at least six generations before use in experiments.

### Bone marrow derived macrophages culture

Bone marrow cells were collected by flushing the femurs and tibias of 8 to 10-week-old C57BL/6 mice and passed through filter. Collected cells were treated with ACK lysis buffer and subsequently washed with PBS. Cells were cultured in the DMEM media supplemented with 10% FBS, 1X antibiotic and 20 % L929 conditioned media for 6-8 days.

### Flow cytometry analysis of cell surface receptors

Single cell suspensions were made from peritoneal macrophages and bone marrow derived macrophages. After blocking with CD16/CD32 antibody, cells were stained with respective antibodies at 4° C for 30 min. Subsequently, cells were washed multiple times with the washing buffer (5 % FBS in 1X PBS), resuspended in the buffer and observed in BD LSRFortessa^™^. The analysis was done by using FlowJo software.

### Flow cytometry - Intracellular staining

For Ki67 and γH2A.X staining, cell surface markers-stained cells were resuspended in BD cytofix/cytoperm buffer and incubated at 4°C for 30 minutes. Subsequently cells were washed with 1X BD perm/wash buffer. Washed cells were resuspended in antibody containing 1X BD perm/wash buffer and incubated for 20 mins at room temperature. Fixed and permeabilized cells were further washed and resuspended in perm/wash buffer and staining buffer respectively. Stained cells were analyzed by LSR Fortessa (BD Biosciences).

For phosphorylated ATM, ATR and p53, eBioscience™ Foxp3 / Transcription Factor Staining kit was used as per manufacture’s protocol.

### Annexin V - PI assay

Cells were harvested and washed with cold PBS. Further cells were resuspended in binding buffer (0.01M HEPES pH-7.4, 0.14M NaCl, 2.5 mM CaCl_2_) at a concentration of 10^6^ cells per ml. Cells were further aliquoted as 100 μl (10^5^ cells) per tube. 5μl of FITC Annexin V and 2.5 μl of PI (1mg/ml) were added in the tubes. After gentle vortexing, cells were incubated at room temperature for 15 min in dark. Cells were further diluted in the binding buffer and flow cytometry analysis was performed using LSR Fortessa (BD Biosciences). Data were analyzed using FlowJo software.

### BrdU based cell proliferation assay

For identification of actively proliferating cells, cells were exposed to BrdU for longer hours. Cells were incubated with 50 μg/ml BrdU for 15 hrs. and 24 hrs. for earlier time points (day3 and day5) and day7 respectively. After incubation, cells were washed and stained with cell surface markers. Stained cells were fixed and permeabilized using BD Pharmingen™ APC BrdU Kit. Subsequently cells were stained with BrdU antibody and analyzed by flow cytometry.

### PI staining

Cells were fixed with 70% ethanol for two hours at 4°C and subsequently washed with PBS. RNase A treatment was given at 37°C for 30 mins. After washing the fixed cells with PBS, cells were resuspended in PI containing PBS and incubated for 30 min at room temperature. Cells were analyzed by flow cytometry.

### Antibodies

For western H3.3 antibody (Sigma Aldrich, WH0003021M1), H3 antibody (Abcam, ab1791), TFIIB antibody (Abcam, ab109518), β-Actin antibody (Cell Signaling Technology, 3700), Stat1 antibody (BD Biosciences, 610115), pY701 Stat1 antibody (BD Biosciences, 612232), Parp1 antibody (Cell Signaling Technology, 9542), p53 antibody (Leica Biosystems, NCL-L-p53-CM5p), p21 antibody (Santa Cruz Biotechnology, sc6246), ATM antibody (Abcam, ab78), ATR antibody (Santa Cruz Biotechnology, sc515173), Caspase7 (Cell Signaling Technology, 9492), Caspase9 (Cell Signaling Technology, 9508), Caspase3 antibody (Cell Signaling Technology, 9662) were used.

For CUT & RUN sequencing, HA antibody (Abcam, ab9110), H3K27ac antibody (Abcam, ab4729), H3K36me3 antibody (Abcam, ab9050) were used.

For flow cytometry CD16/CD32 antibody (Biolegend, 101320), CD11b antibody (BD, 562605), F4/80 antibody (Biolegend,123116), Brilliant Violet 421™ anti-mouse/human CD11b antibody (Biolegend, 101235), FITC anti-mouse/human CD11b antibody (Biolegend, **101206),** APC anti-mouse F4/80 antibody (Biolegend, 123115), PE/Cyanine7 anti-mouse F4/80 antibody (Biolegend, 123113), APC anti-mouse Ki-67 antibody (Biolegend, 652405), BD Horizon™ BV421 Mouse Anti-H2AX (pS139) (BD, 564720), BD Horizon™ BV421 Mouse IgG1, k Isotype Control (BD, 562438), Phospho-ATM (Ser1981) monoclonal antibody, eFluor™ 660 (Thermo Fisher Scientific, 50-9046-42), ATR (phospho Thr1989) antibody (GeneTex, GTX128145), phospho-p53 (Ser15) antibody (Cell Signaling Technology, 9284), AF633 (were used.

For IFNAR1 neutralization, MAR1-5A3 antibody (Affymetrix eBioscience, 16-5945-85), IgG1 κ isotype control (eBioscience, 14-4714-85) were used.

### Quantitative real time PCR, primer list

Total RNA was isolated using Trizol. cDNA was synthesized using High-capacity cDNA reverse transcription kit (Applied Biosystems, Thermo Fisher Scientific). Quantitative real time PCR was performed using Fast SYBR^™^ Green Master Mix (Applied Biosystems, Thermo Fisher Scientific). Quantitative differences were calculated using the 2 CT method relative to *Gtf2b*.

### Blocking of IFNAR1 receptor

WT and cKO BM cells were incubated with F12/DMEM supplemented with 20% L929 conditioned medium. IFNAR1 monoclonal antibody or IgG1 κ isotype control antibodies were added at a concentration of 10 μg/mL at 37 °C. Antibodies were added since day0 and media was supplemented with fresh antibodies every 12 hrs. On Day 6, the media containing antibodies were removed, and cells were lysed with trizol.

### RNA seq and data analysis

For BMDM, total RNA was prepared using Quick-RNA Mini prep Kit (Zymo Research). RNA integrity was confirmed by running the RNA sample on an Agilent Bioanalyzer RNA 6000 Nano chip to determine the RNA integrity number. mRNA enrichment was done using NEBNext® rRNA depletion kit. RNA seq libraries were prepared using NEBNext® Ultra^™^ II RNA library prep kit.

Peritoneal macrophages were isolated using miltney kit. Total RNA was purified by trizol extraction. Library was prepared using SMART-Seq v4 Ultra Low Input RNA Kit (Takara bio USA, CA) with 7-cycle amplification.

Samples were pooled and sequenced (paired end) on the Illumina NextSeq 500 and NextSeq2000.

For data analysis, quantification, and quality-control pipeline (CCBR pipeliner v4.0.6) was used. Briefly, adapter sequences are removed using Cutadapt and paired-end reads were aligned to the *Mus musculus* reference genome mm10 using STAR aligner. Gene and isoform expression levels were quantified using RSEM. For differential expression analysis SARTools1.8.1 [50] were used with q value of <0.05. Heatmaps were depicted with Morpheus (https://software.broadinstitute.org/morpheus).

Genes with baseMean > 20 (RNA seq) were considered expressed genes (n=12922). Remaining genes and isoforms were considered as silent genes. Cell cycle gene list was used as described [18]. To define ISGs, we stimulated WT BMDMs with interferon γ for different time points (2,4,6,8,10 and 12 hrs.) and performed RNA seq. All the upregulated genes (merged and duplicates were removed) were defined as ISGs.

All known mouse genes were downloaded from UCSC table browser.

### CUT & RUN seq and data analysis

CUT & RUN was performed as reported earlier [51]. Cells were collected and washed twice with the buffer (20 mM HEPES pH 7.5, 150 mM NaCl, 0.5 mM Spermidine, 1x Protease inhibitor cocktail). Concanavalin A-coated magnetic beads were washed twice with the binding buffer (20 mM HEPES-KOH pH-7.9, 10 mM KCl, 1 mM CaCl2,1 mM MnCl2). Cells were resuspended in wash buffer, mixed with the bead suspension. The cell-bead mix was rotated for 10 min at room temperature, placed on magnetic rack and liquid was discarded. Beads were further dislodged in antibody buffer (wash buffer containing 2mM EDTA, 0.02% digitonin and respective antibody). Antibody incubation was done overnight at 4 ^0^C. After incubation, beads were placed on magnetic rack and all the liquid was removed. Subsequently, beads were washed with digitonin buffer (wash buffer containing 0.02 % digitonin). Further, beads were resuspended in pA-MNase (700ng/ml) containing digitonin buffer and incubated on tube rotator for 1 hour at 4^0^ C. After 1 hour, beads were washed twice with digitonin buffer, resuspended in 100 µl of digitonin buffer, and placed on a cold block to reach 0 ^0^C. 3 µl of 100 mM CaCl2 was mixed in the bead suspension which was further incubated at 4 ^0^C for 30’. The reaction was stopped by adding 100 µl of 2X stop buffer (0.34 M NaCl, 20 mM EDTA, 4 mM EGTA, 0.02 % digitonin, 50 μg/ml RNase A, 50 μg/ml glycogen, 20 pg/ml yeast spike in DNA). To release the fragments, beads were incubated at 37^0^ C for 10’, centrifuged at 4 ^0^C for 5’. The supernatant was collected from the tubes on the magnetic rack. For histone antibodies, DNA was extracted by using Qiagen spin column, while in the case of transcription factors, PCI based extraction was done after adding 0.1 % SDS and 150 μg/ml of proteinase K in the sample and incubating it at 70^0^ C for 10‘. (Spike in) CUT & RUN seq libraries were prepared using NEBNext® Ultra^™^ II DNA library kit (Illumina). Samples were pooled and paired end sequencing (25 bp) was performed on the Illumina NextSeq 500.

For data analysis, CUT & RUN -seq reads were mapped to the mouse reference genome (mm10) using Bowtie2 v.2.3.4.1 with the following parameters: --local --very-sensitive-local --no-unal --no-mixed --no-discordant --phred33 -I 10 -X 700. The peak calling was performed using homer. The bam files were converted to normalized bigWig files by using scale factor, deeptools bamCoverage command[52]. DiffBind was used to analyze differential binding[53]. We used GREAT to further identify the genes associated with the peaks[54].

### ATAC seq and data analysis

ATAC seq was performed as described previously [55, 56]. Briefly, 50,000 cells were lysed to prepare crude nuclei extract. The nuclei were resuspended in transposition reaction mix and incubated at 37^0^ C for 30’. Transposed DNA was collected using DNA Clean & Concentrator™-5 purification columns. Eluted DNA was amplified using NEBNext® Ultra^™^ II Q5 Mater Mix and size selected (Negative and positive selection). Purified DNA samples were assessed using Bioanalyzer High sensitivity DNA analysis kit. Samples were subsequently pooled and sequenced using paired-end sequencing on the Illumina NextSeq 500. Paired reads were aligned to mouse reference genome mm10 using Bowtie2-2.5.1 Peak calling was performed by using Genrich with default settings. To identify differential ATAC-seq peaks between WT and cKO samples, R package DiffBind was used. MA plot, cluster correlation analysis plots were generated using DiffBind package. For motif analysis, homer was used.

### MNase digestion

Day7 BMDMs were washed with ice cold PBS and resuspended in buffer (50mM NaCl, 10mM Tris-Cl, pH-7.5, 5 mM MgCl2, 1mM CaCl2, 0.2 % NP-40, 1X protease inhibitor cocktail). After the lysis, nuclei were resuspended in buffer and incubated with 1U MNase for indicated time points (5, 10, 15, 20, 25 and 30 mins). The enzymatic reaction was stopped by addition of EGTA. After phenol, chloroform, isoamyl alcohol treatment of the reaction, aqueous phase DNA was collected and further purified by Qiagen purification kit.

### Acidic extraction of histones

Acidic extraction of histones was performed as reported (https://www.abcam.com/protocols/histone-extraction-protocol-for-western-blot). Briefly, WT and KO cells were washed with ice cold PBS. Subsequently cells were lysed in triton extraction buffer (PBS containing 0.5 % Triton X 100 (v/v), 2 mM phenylmethylsulphonyl fluoride (PMSF), 0.02 % (w/v) NaN3) for 10 min on ice. Nuclei pellets were collected by centrifugation at 4 ^0^C for 10 min at 6500 g. Nuclei were further washed in triton extraction buffer. Washed nuclei were resuspended in of 0.2N HCl and acid extracted overnight at 4 ^0^C. Histones (supernatant) were collected by centrifugation at 4 ^0^C for 10 min at 6500 g. Acid extracted histones were further neutralized with 2M NaOH.

### Whole cell extract and nuclear extract preparation

For cell lysates preparation, cells were lysed in Radio Immunoprecipitation Assay (RIPA) buffer (25mM Tris-HCl, pH7.6, 150mM NaCl (sodium chloride), 1% NP-40, 0.1% SDS (sodium dodecyl sulfate), 0.1% sodium deoxycholate) with protease and phosphatase inhibitors for 1h at 4°C. Whole cell lysates were clarified by centrifugation at 12,000 rpm for 10min at 4°C. Nuclear extracts were prepared as described [57].

### Western blotting

Cell lysates and histones were separated on NuPAGE^TM^4-12 % and 12% Bis-Tris gel respectively. Proteins were subsequently transferred to PVDF membrane (Millipore). Blocking was performed at room temperature for 1 hour with 2.5% BSA. Blots were incubated overnight with primary antibody at 4 ^0^C and then washed twice for 10 min each. Secondary antibody incubation was performed at room temperature for 1 hour. To remove nonspecific binding, blots were washed again and then incubated with ECL for 5 min and analyzed with Azure c600 Biosystems.

### Data availability

All high-throughput sequence datasets generated in this paper are available at GEO accession nos. GSE285139, GSE285140 and GSE285141.

## Acknowledgements

We acknowledge NICHD animal facility staff for caring for our mice and for other technical assistance. We gratefully acknowledge colleagues in our lab and elsewhere for technical advice and valuable discussions. This work was supported by the NICHD Intramural programs ZIA HD008015-13.

## References

1. Ng, R.K. and J.B. Gurdon, Epigenetic memory of an active gene state depends on histone H3.3 incorporation into chromatin in the absence of transcription. Nat Cell Biol, 2008. 10(1): p. 102–9.

2. Wong, L.H., et al., Histone H3.3 incorporation provides a unique and functionally essential telomeric chromatin in embryonic stem cells. Genome Res, 2009. 19(3): p. 404–14.

3. Goldberg, A.D., et al., Distinct factors control histone variant H3.3 localization at specific genomic regions. Cell, 2010. 140(5): p. 678–91.

4. Jang, C.W., et al., Histone H3.3 maintains genome integrity during mammalian development. Genes Dev, 2015. 29(13): p. 1377–92.

5. Guo, P., et al., Histone variant H3.3 maintains adult haematopoietic stem cell homeostasis by enforcing chromatin adaptability. Nat Cell Biol, 2022. 24(1): p. 99–111.

6. Funk, O.H., et al., Postmitotic accumulation of histone variant H3.3 in new cortical neurons establishes neuronal chromatin, transcriptome, and identity. Proc Natl Acad Sci U S A, 2022. 119(32): p. e2116956119.

7. Saito, Y., et al., Plasma cell differentiation is regulated by the expression of histone variant H3.3. Nat Commun, 2024. 15(1): p. 5004.

8. Li, P., et al., Cytochrome c and dATP-dependent formation of Apaf-1/caspase-9 complex initiates an apoptotic protease cascade. Cell, 1997. 91(4): p. 479–89.

9. Kaufmann, S.H., et al., Specific proteolytic cleavage of poly(ADP-ribose) polymerase: an early marker of chemotherapy-induced apoptosis. Cancer Res, 1993. 53(17): p. 3976–85.

10. Shiloh, Y. and Y. Ziv, The ATM protein kinase: regulating the cellular response to genotoxic stress, and more. Nat Rev Mol Cell Biol, 2013. 14(4): p. 197–210.

11. Zou, L. and S.J. Elledge, Sensing DNA damage through ATRIP recognition of RPA-ssDNA complexes. Science, 2003. 300(5625): p. 1542–8.

12. Bonner, W.M., et al., GammaH2AX and cancer. Nat Rev Cancer, 2008. 8(12): p. 957–67.

13. Ciccia, A. and S.J. Elledge, The DNA damage response: making it safe to play with knives. Mol Cell, 2010. 40(2): p. 179–204.

14. Zeman, M.K. and K.A. Cimprich, Causes and consequences of replication stress. Nat Cell Biol, 2014. 16(1): p. 2–9.

15. Groelly, F.J., et al., Targeting DNA damage response pathways in cancer. Nat Rev Cancer, 2023. 23(2): p. 78–94.

16. Bunz, F., et al., Requirement for p53 and p21 to sustain G2 arrest after DNA damage. Science, 1998. 282(5393): p. 1497–501.

17. Vogelstein, B., D. Lane, and A.J. Levine, Surfing the p53 network. Nature, 2000. 408(6810): p. 307–10.

18. Wu, T., et al., Bromodomain protein BRD4 directs mitotic cell division of mouse fibroblasts by inhibiting DNA damage. iScience, 2024. 27(7): p. 109797.

19. Diehl, F.F., et al., Nucleotide imbalance decouples cell growth from cell proliferation. Nat Cell Biol, 2022. 24(8): p. 1252–1264.

20. Lane, A.N. and T.W. Fan, Regulation of mammalian nucleotide metabolism and biosynthesis. Nucleic Acids Res, 2015. 43(4): p. 2466–85.

21. Brzostek-Racine, S., et al., The DNA damage response induces IFN. J Immunol, 2011. 187(10): p. 5336–45.

22. Mboko, W.P., et al., Coordinate regulation of DNA damage and type I interferon responses imposes an antiviral state that attenuates mouse gammaherpesvirus type 68 replication in primary macrophages. J Virol, 2012. 86(12): p. 6899–912.

23. Yu, Q., et al., DNA-damage-induced type I interferon promotes senescence and inhibits stem cell function. Cell Rep, 2015. 11(5): p. 785–797.

24. Walter, D., et al., Exit from dormancy provokes DNA-damage-induced attrition in haematopoietic stem cells. Nature, 2015. 520(7548): p. 549–52.

25. Li, D. and M. Wu, Pattern recognition receptors in health and diseases. Signal Transduct Target Ther, 2021. 6(1): p. 291.

26. Akira, S. and K. Takeda, Toll-like receptor signalling. Nat Rev Immunol, 2004. 4(7): p. 499–511.

27. Decout, A., et al., The cGAS-STING pathway as a therapeutic target in inflammatory diseases. Nat Rev Immunol, 2021. 21(9): p. 548–569.

28. Seth, R.B., et al., Identification and characterization of MAVS, a mitochondrial antiviral signaling protein that activates NF-kappaB and IRF 3. Cell, 2005. 122(5): p. 669–82.

29. Thoresen, D., et al., The molecular mechanism of RIG-I activation and signaling. Immunol Rev, 2021. 304(1): p. 154–168.

30. Kim, T.K., et al., Chemotherapeutic DNA-damaging drugs activate interferon regulatory factor-7 by the mitogen-activated protein kinase kinase-4-cJun NH2-terminal kinase pathway. Cancer Res, 2000. 60(5): p. 1153–6.

31. Pan, D., et al., SETDB1 Restrains Endogenous Retrovirus Expression and Antitumor Immunity during Radiotherapy. Cancer Res, 2022. 82(15): p. 2748–2760.

32. Gehre, M., et al., Lysine 4 of histone H3.3 is required for embryonic stem cell differentiation, histone enrichment at regulatory regions and transcription accuracy. Nat Genet, 2020. 52(3): p. 273–282.

33. Martire, S., et al., Phosphorylation of histone H3.3 at serine 31 promotes p300 activity and enhancer acetylation. Nat Genet, 2019. 51(6): p. 941–946.

34. Tafessu, A., et al., H3.3 contributes to chromatin accessibility and transcription factor binding at promoter-proximal regulatory elements in embryonic stem cells. Genome Biol, 2023. 24(1): p. 25.

35. Trovato, M., et al., Histone H3.3 lysine 9 and 27 control repressive chromatin at cryptic enhancers and bivalent promoters. Nat Commun, 2024. 15(1): p. 7557.

36. Bachu, M., et al., A versatile mouse model of epitope-tagged histone H3.3 to study epigenome dynamics. J Biol Chem, 2019. 294(6): p. 1904–1914.

37. McKittrick, E., et al., Histone H3.3 is enriched in covalent modifications associated with active chromatin. Proc Natl Acad Sci U S A, 2004. 101(6): p. 1525–30.

38. Hake, S.B., et al., Expression patterns and post-translational modifications associated with mammalian histone H3 variants. J Biol Chem, 2006. 281(1): p. 559–68.

39. Hnisz, D., et al., Super-enhancers in the control of cell identity and disease. Cell, 2013. 155(4): p. 934–47.

40. Sun, Z., et al., H3K36me3, message from chromatin to DNA damage repair. Cell Biosci, 2020. 10: p. 9.

41. Chantalat, S., et al., Histone H3 trimethylation at lysine 36 is associated with constitutive and facultative heterochromatin. Genome Res, 2011. 21(9): p. 1426–37.

42. el-Deiry, W.S., et al., WAF1, a potential mediator of p53 tumor suppression. Cell, 1993. 75(4): p. 817–25.

43. el-Deiry, W.S., et al., WAF1/CIP1 is induced in p53-mediated G1 arrest and apoptosis. Cancer Res, 1994. 54(5): p. 1169–74.

44. Banaszynski, L.A., et al., Hira-dependent histone H3.3 deposition facilitates PRC2 recruitment at developmental loci in ES cells. Cell, 2013. 155(1): p. 107–20.

45. Saxena, S. and L. Zou, Hallmarks of DNA replication stress. Mol Cell, 2022. 82(12): p. 2298–2314.

46. Herr, L.M., et al., Replication stress as a driver of cellular senescence and aging. Commun Biol, 2024. 7(1): p. 616.

47. Honda, K., et al., IRF-7 is the master regulator of type-I interferon-dependent immune responses. Nature, 2005. 434(7034): p. 772–7.

48. Wiedemann, S.M., et al., Identification and characterization of two novel primate-specific histone H3 variants, H3.X and H3.Y. J Cell Biol, 2010. 190(5): p. 777–91.

49. Ishikawa, H. and G.N. Barber, STING is an endoplasmic reticulum adaptor that facilitates innate immune signalling. Nature, 2008. 455(7213): p. 674–8.

50. Varet, H., et al., SARTools: A DESeq2- and EdgeR-Based R Pipeline for Comprehensive Differential Analysis of RNA-Seq Data. PLoS One, 2016. 11(6): p. e0157022.

51. Skene, P.J., J.G. Henikoff, and S. Henikoff, Targeted in situ genome-wide profiling with high efficiency for low cell numbers. Nat Protoc, 2018. 13(5): p. 1006–1019.

52. Ramirez, F., et al., deepTools2: a next generation web server for deep-sequencing data analysis. Nucleic Acids Res, 2016. 44(W1): p. W160–5.

53. Ross-Innes, C.S., et al., Differential oestrogen receptor binding is associated with clinical outcome in breast cancer. Nature, 2012. 481(7381): p. 389–93.

54. McLean, C.Y., et al., GREAT improves functional interpretation of cis-regulatory regions. Nat Biotechnol, 2010. 28(5): p. 495–501.

55. Buenrostro, J.D., et al., Transposition of native chromatin for fast and sensitive epigenomic profiling of open chromatin, DNA-binding proteins and nucleosome position. Nat Methods, 2013. 10(12): p. 1213–8.

56. Buenrostro, J.D., et al., ATAC-seq: A Method for Assaying Chromatin Accessibility Genome-Wide. Curr Protoc Mol Biol, 2015. **109**: p. 21 29 1-21 29 9.

57. Abmayr, S.M., et al., Preparation of nuclear and cytoplasmic extracts from mammalian cells. Curr Protoc Mol Biol, 2006. Chapter 12: p. Unit 12 1.

