## Supplementary data for "Histone H3.3 ensures cell proliferation and genomic stability during myeloid cell development"

### Supplementary Figure legends

Supplementary Fig 1. Recombination and LysMcre based conditional deletion of histone H3.3 genes

a) Schematic representation of *H3f3a* and b) *H3f3*b alleles, created for this study.

c) Recombined and cre excised bands of *H3f3a* and *H3f3b* genes by LysMcre.

a) qRTPCR confirmation of *H3.3* deletion at RNA level in BMDMs and peritoneal macrophages

b) Western blot confirmation of H3.3 deletion at protein level.

c) Gene tracks show *H3f3a* and *H3f3b* gene deletion in cKO cells.

d) Representative flow cytometry plots for peritoneal macrophages upon staining with CD11b and F4/80 markers. Total number of peritoneal macrophages (right) were obtained from WT and cKO mice.

*Note: H3f3b gene deletion was found to be fluctuating as we observed complete deletion as well as partial deletion of the gene in different cKO samples.*

Supplementary Fig 2. Surface marker expression, BrdU incorporation schematic and H3.3 deletion on day3, 5 and day7 in WT and cKO cells

a) Representative flow cytometry plots for CD11b and F4/80 expression in WT and cKO cells on day3, 5 and day7.

b) Normalized read counts of *H3f3a* gene in WT and cKO cells at different time points.

c) Normalized read counts of *H3f3b* gene in WT and cKO cells at different time points.

*Note: H3f3b gene deletion was found to be fluctuating as we observed complete deletion as well as partial deletion of the gene in different KO samples.*

d) Schematic representation of BrdU incorporation in WT and cKO cells.

Supplementary Fig 3. Mki67 expression and apoptosis in cKO cells.

a) Graph shows difference in average read counts (normalized) of Mki67 between WT and KO cells

b) IGV browser screenshot shows lower expression of Mki67 RNA on day3.

c) Ki67 protein expression on day3,5 and 7.

d) % BMDM represent the average of three WT and three *H3.3* cKO mice ± SD. P values were calculated using unpaired t test.
e) Cleavage of Parp1 and caspase genes is shown through immunoblotting.

Supplementary Fig 4. ISG upregulation in cKO cells is dependent on interferon feedback.

a) Schematic representation of IFNAR assay
b) Heatmap shows normalized read counts for KO upregulated ISGs for WT and cKO samples for all three conditions (No antibody, control antibody and IFNAR1 antibody).
c) Gene tracks of Ifi204 and Stat1 show decrease in RNA expression upon IFNAR neutralization.
d) qRTPCR based confirmation of IFNAR assay by using *Ifi204* and *Stat1* genes. n=3 mice. P values were calculated using ordinary one way ANOVA test. The error bars represent standard error of mean. NS - Not significant

Supplementary Fig 5 *H3.3* deletion does not hinder M-CSF induced BMDM differentiation.

a) IGV gene tracks show equal expression of lineage determining transcription factors (*Spi1* and *Irf8*) in WT and cKO BMDMs.
b) Genic expression of macrophage marker gene *Adgre1/F4/80* and *Itgam/CD11b*.
c) Representative flow cytometry plots for staining of CD11b and F4/80 protein in WT and cKO cells.
d) MFI values were calculated for F4/80 and CD11b protein expression in WT and cKO cells
e) Heatmaps depict upregulation of ISGs between WT vs cKO BMDMs, WT vs IFN$\gamma$ treated (12hrs) WT BMDMs and KO vs IFN$\gamma$ treated (12hrs) KO BMDMs.
f) IGV gene tracks show increased genic expression of *Ifi44* and *Oasl1* in WT vs cKO BMDMs, WT vs IFN$\gamma$ treated (12hrs) WT BMDMs and cKO vs IFN$\gamma$ treated (12hrs) cKO BMDMs.

Supplementary Fig 6 H3.3 distribution and differential chromatin accessibility in WT and cKO cells

a) Gene tracks showing deposition of H3.3 on expressed gene Itgam,
b) Gene tracks showing deposition of H3.3 on cell cycle genes Kif5b and E2f3.
c) GO analysis of increased and decreased accessible peaks in cKO cells.
d) Volcano plot showing a correlation between ATAC seq peaks and RNA seq data. The nearest genes for increased or decreased ATAC peaks correlate with genes up or downregulated in H3.3 cKO BMDM.
e) Motif analysis of KO gained, or KO lost peaks is shown.

Supplementary Fig 7. Differential binding analysis of H3K27ac and H3K36me3

a) H3K27ac deposition on gene bodies +3kb/-3kb in WT cells and
b) its genome wide distribution
c) MA plot represents increased or decreased H3K27ac peaks upon conditional deletion of *H3.3*
d) GO analysis of KO gained and lost peaks of H3K27ac
e) H3K36me3 distribution on gene bodies and
a) its distribution across genome in WT cells
b) MA plot for H3K36me differential binding in WT and cKO BMDMs
c) H3.3, H3K27ac and H3K36me3 distribution across known mouse genes
d) H3.3, H3K27ac and H3K36me3 distribution on macrophage marker genes *Itgam*, *Adgre1*.

Supplementary Fig1

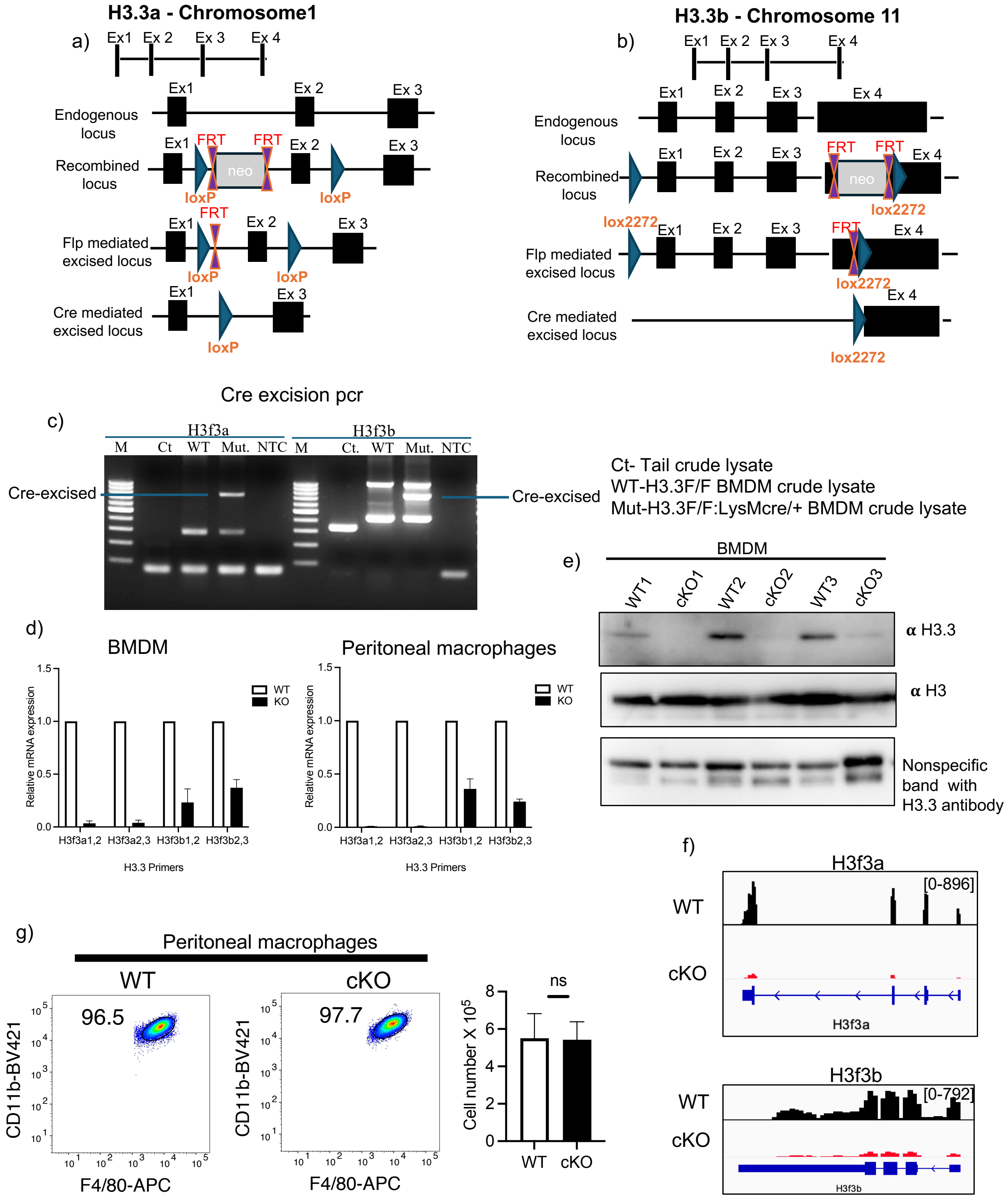

Supplementary Fig2

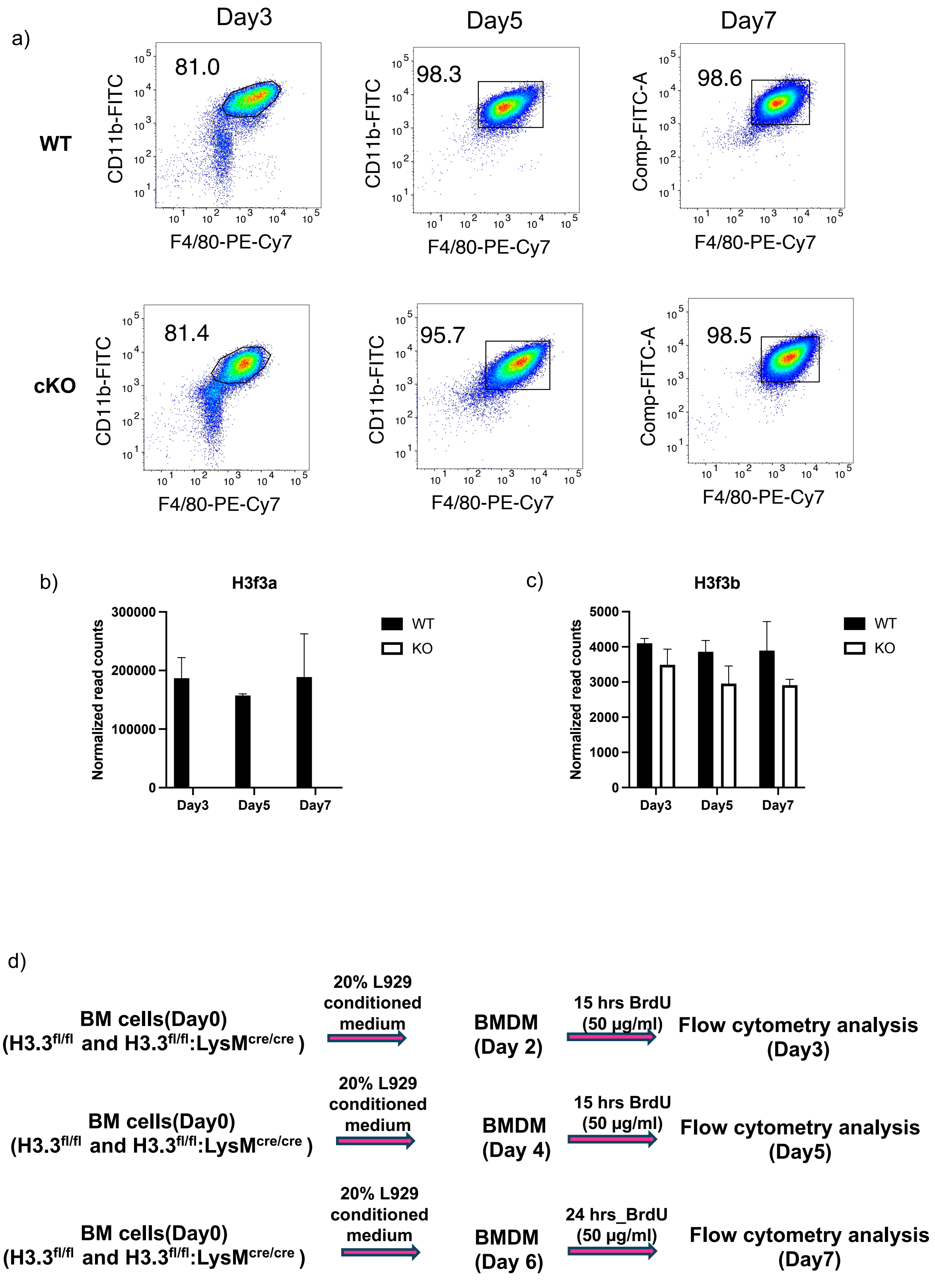

Supplementary Fig3

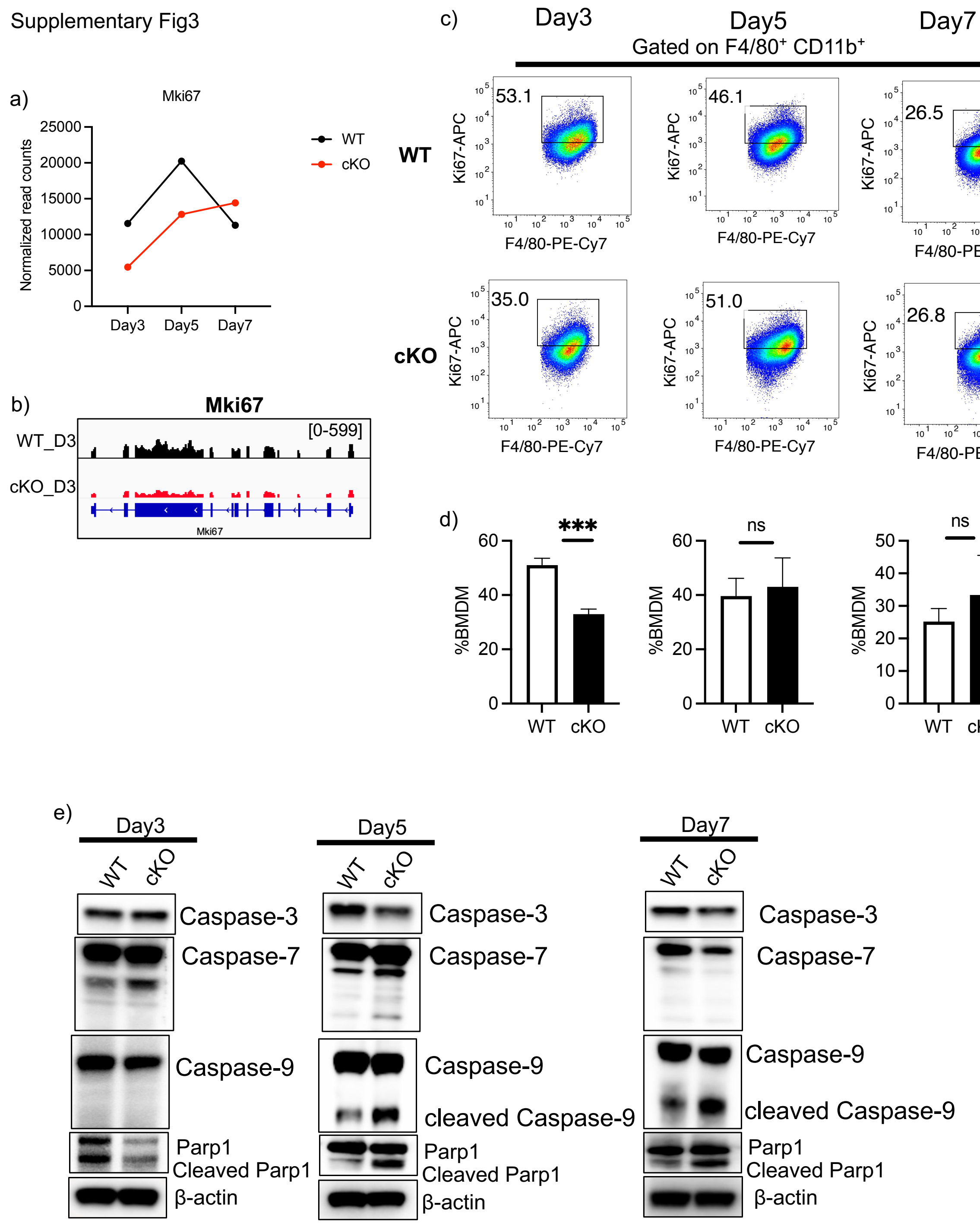

Supplementary Fig4

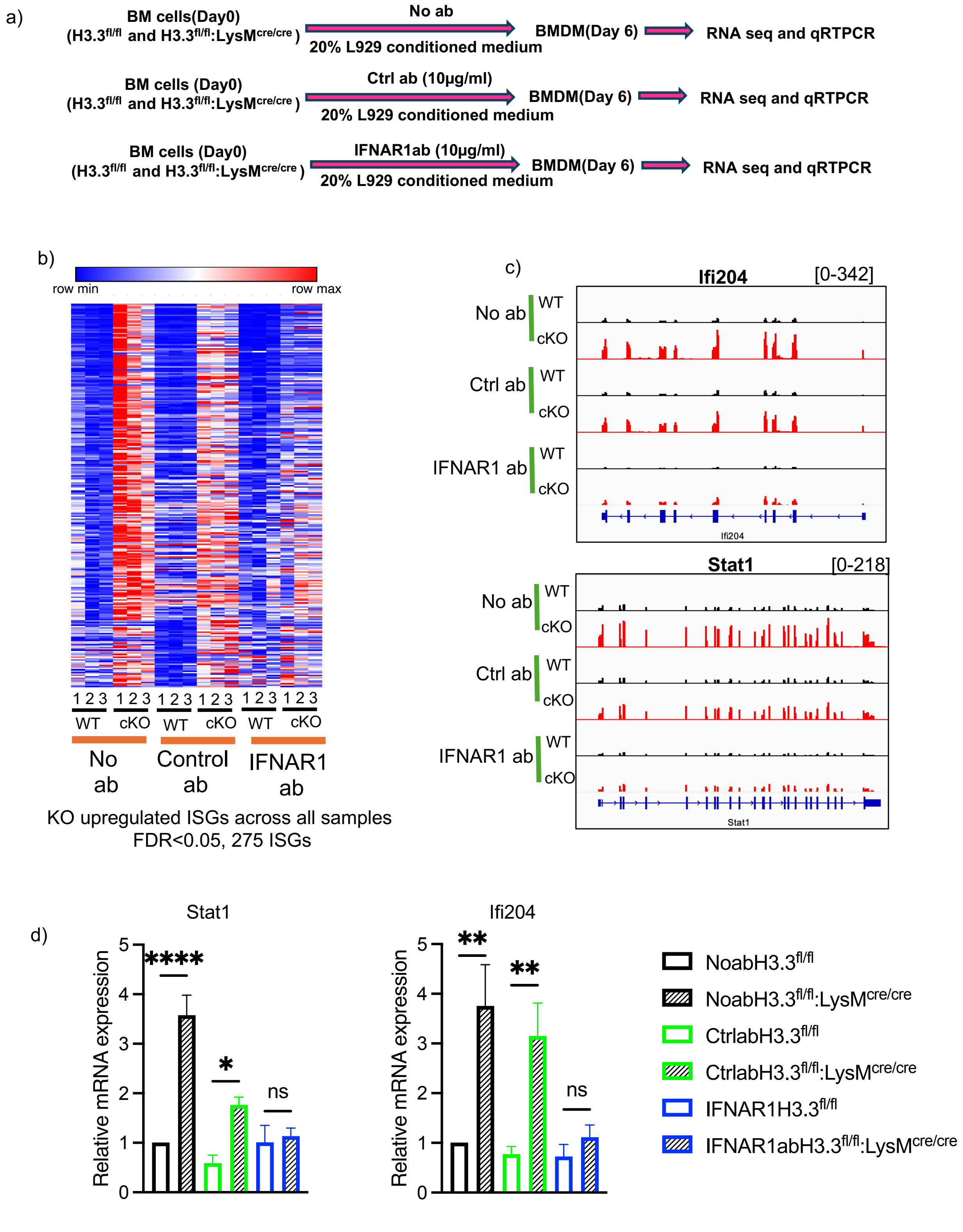

Supplementary Fig5

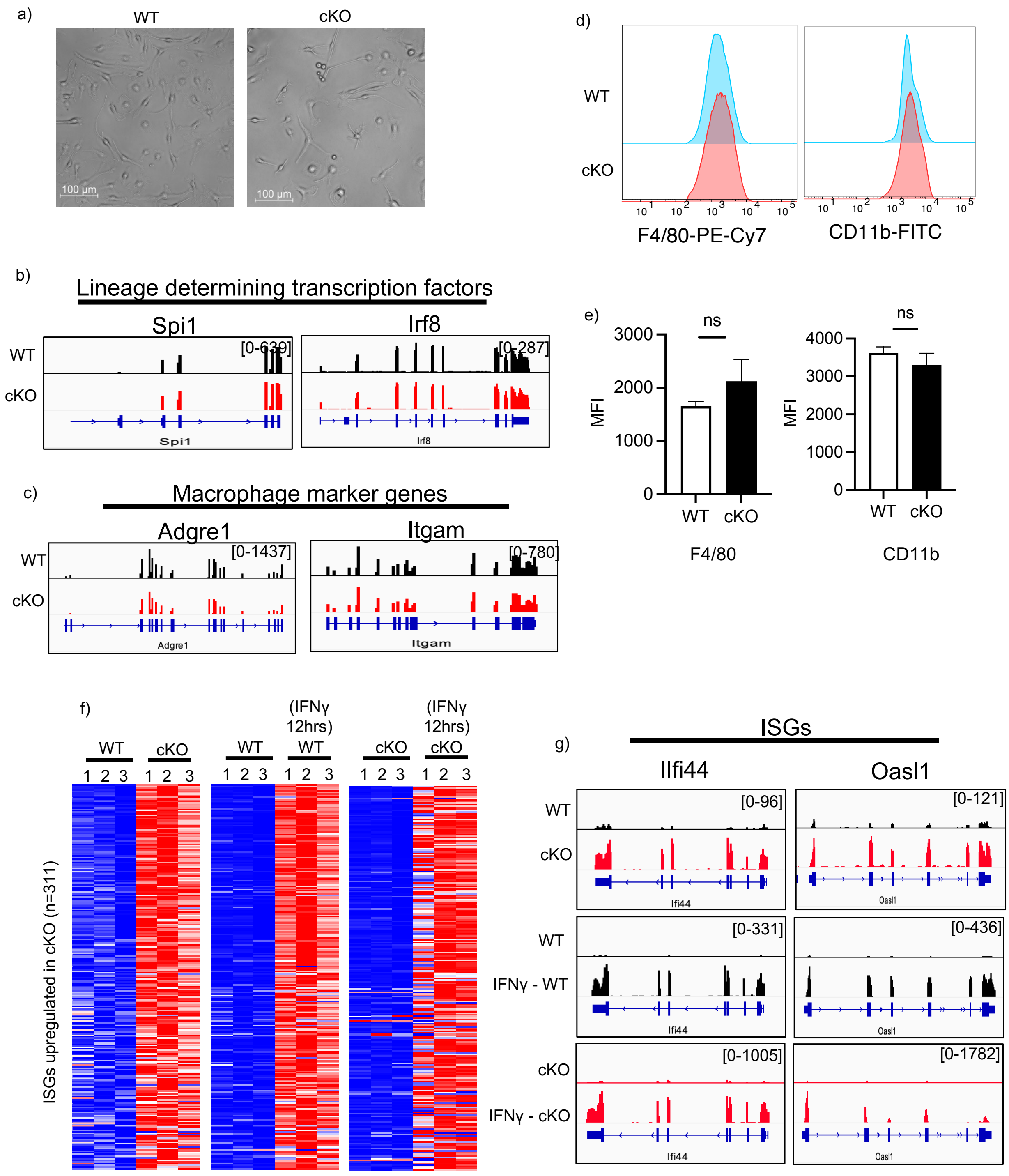

Supplementary Fig6

a) b)

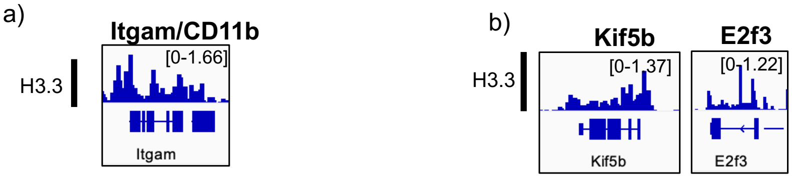


c)

| | GO terms | BinomP |
|---|---|---|
| Increased accessibility peaks | defense response to virus | 5.86E-15 |
| | positive regulation of apoptotic process | 2.99E-08 |
| | ribonucleoprotein complex biogenesis | 4.02E-08 |
| | cellular response to DNA damage stimulus | 5.14E-07 |
| | regulation of G1/S transition of mitotic cell cycle | 4.85E-06 |
| | pyrimidine ribonucleoside catabolic process | 1.27E-05 |
| Decreased accessibility peaks | regulation of cell adhesion | 1.37E-30 |
| | regulation of leukocyte activation | 2.96E-27 |
| | positive regulation of immune system process | 7.32E-27 |
| | regulation of cell activation | 4.31E-26 |
| | vesicle-mediated transport | 4.60E-25 |
| | response to external biotic stimulus | 2.08E-23 |
| | lipid biosynthetic process | 6.60E-10 |

d)

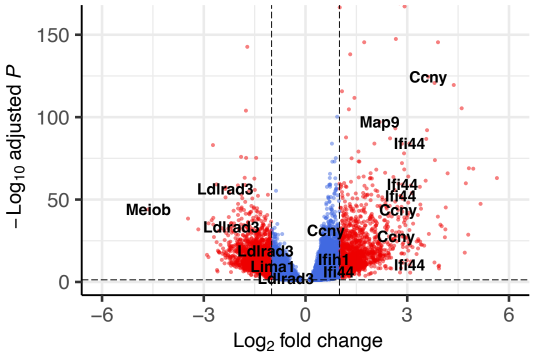


e)

cKO gained motifs

| Rank | Motif | P-value | T/B(%) | TF |
|---|---|---|---|---|
| 1 | 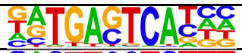 | 1e-286 | 36.02/8.28 | Atf3(bZIP)(0.990) |
| 2 | 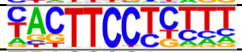 | 1e-161 | 37.02/13.82 | Sfpi(0.953) |
| 3 | 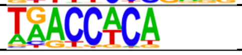 | 1e-61 | 31.42/16.94 | Runx(0.914) |

cKO lost motifs

| Rank | Motif | P-value | T/B(%) | TF |
|---|---|---|---|---|
| 1 | 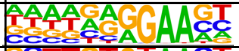 | 1e-623 | 67.11/20.58 | Sfpi (0.906) |
| 2 | 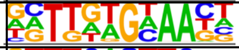 | 1e-81 | 37.71/21.86 | CEBPD (0.887) |
| 3 | 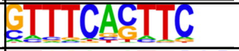 | 1e-74 | 8.58/2.08 | Pu.1-IRF (0.943) |
| 4 | 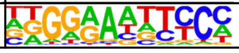 | 1e-73 | 13.79/4.89 | NFkB-p65 (0.893) |

Supplementary Fig7

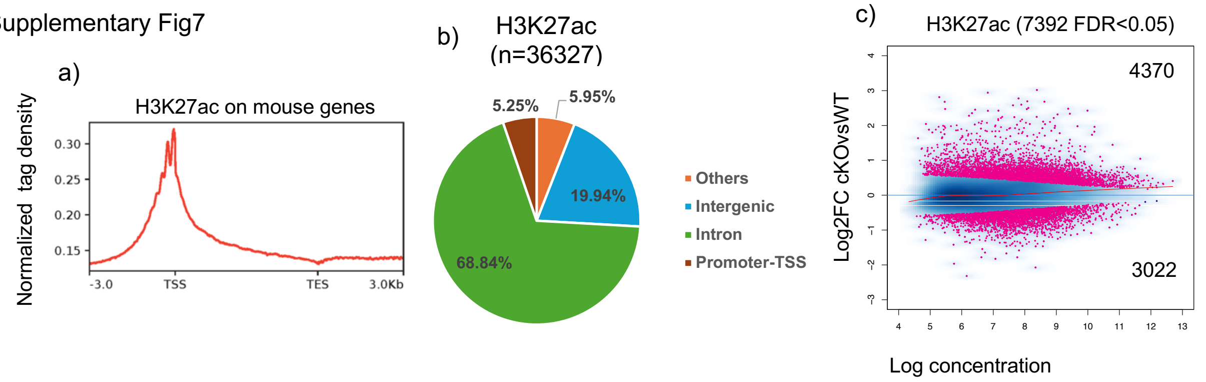


d)

| | GO terms | BinomP |
|---|---|---|
| Increased H3K27ac peaks | defense response to other organism | 3.75E-32 |
| | response to external biotic stimulus | 9.31E-31 |
| | response to virus | 8.28E-24 |
| | cellular response to interferon-beta | 1.94E-17 |
| | positive regulation of programmed cell death | 7.70E-07 |
| Decreased H3K27ac peaks | substrate adhesion-dependent cell spreading | 1.11E-08 |
| | calcium-independent cell-matrix adhesion | 1.36E-06 |
| | blood vessel morphogenesis | 6.00E-06 |
| | actin cytoskeleton reorganization | 3.32E-05 |
| | response to cholesterol | 3.97E-05 |

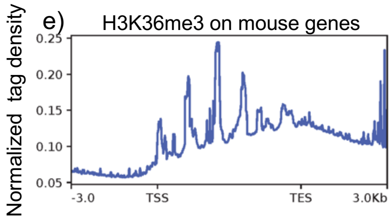


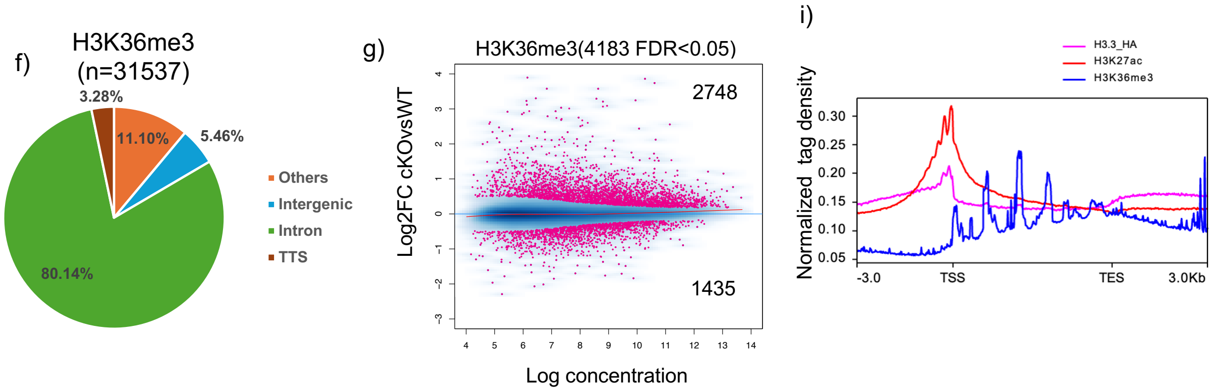


h)

| | GO terms | BinomP |
|---|---|---|
| Increased H3K36me3 peaks | cellular response to interferon-beta | 4.93E-16 |
| | defense response to virus | 7.82E-09 |
| | chromosome organization | 1.13E-07 |
| | regulation of response to DNA damage checkpoint signaling | 1.73E-07 |
| | mitotic nuclear division | 1.94E-07 |
| | mitotic cell cycle | 5.29E-07 |
| Decreased H3K36me3 peaks | lipid biosynthetic process | 3.91E-16 |
| | clustering of voltage-gated sodium channels | 4.75E-16 |
| | sterol biosynthetic process | 1.69E-13 |

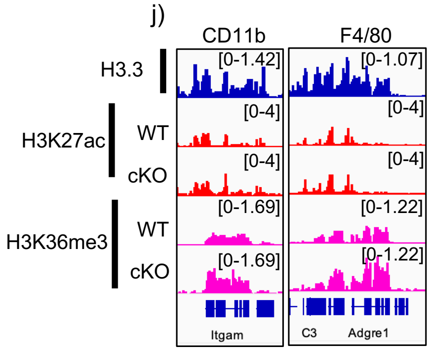

qRT-PCR primers

| Gene Name | FP | RP |
|---|---|---|
| H3f3a (Exon1-2) | CAGCGCCGCCTCTCGCTTG | CAGTCTGCTTTGTACGAGCCATGGTA |
| H3f3a (Exon2-3) | CTACAAAAGCCGCTCGCAAGAGT | TTCTGATAGCGTCTGATTTCACGG |
| H3f3b (Exon1-2) | CGATTGCGGCTCTTGTTCGAG | TTCCCACCGGTGGACTTCCTA |
| H3f3b (Exon2-3) | CCAAGGCGGCTCGGAAAAGC | GGTAACGACGGATCTCTCTCAGA |

Cre excision PCR primers

| | | |
|---|---|---|
| H3f3b | Cre P1 138160 | TGGGATTAAAGAAATGTGTGTGCCAGC |
| | Cre P2 138161 | GGATCCAAAATTCAAGGTCATTCACAGC |
| | Cre P3 138162 | CTCCCCACCACTTGTGAATGTAAAACATTT |
| H3f3a | Cre P1 138090 | CAAACAAGTTTTCAAATGTAGTTATGACTGATGC |
| | Cre P2 138091 | CTAAACATACAAAACGTGGGAACACATTAGC |
| | Cre P3138099 | AAGGGAATCGGATACGTTTCCAATTG |

Flox PCR primers

| | | |
|---|---|---|
| H3f3b | Flp 138154 | AGGAGTGTTAATGCTGTTGCTCTTGTCCTC |
| | Flp 138155 | TCACATCACTGAGGTCTGTGAACAGTCAGT |
| | Flp 138156 | CCTCACCTTGTCGTATTATACTATGCCGATATACT |
| H3f3a | Flp 138094 | CGACCTTTCTGTGTTTGTGGCTTCG |
| | Flp 138095 | GATTTGCGGGCAGTCTGCTTTGTAC |
| | Flp 138097 | CTGTGCTCGACGTTGTCACTGAAGC |
